# Genome-wide chromatin accessibility during early embryogenesis in a generalist cichlid fish

**DOI:** 10.64898/2026.10.30.762802

**Authors:** Midhukrishna, Alan M. Smith, Domino Joyce, Federica Di-Palma, Wilfried Haerty, Tarang K. Mehta

## Abstract

**Background:** Lake Malawi’s haplochromine cichlids are one of the most rapid and species-rich vertebrate adaptive radiations known, generating hundreds of species within < 1 million years, yet how regulatory regions shape their embryonic development remains poorly understood. *Astatotilapia calliptera*, a generalist riverine lineage and proxy for the ancestral stock that seeded this radiation, offers a developmental chromatin baseline against which regulatory divergence in specialised lineages can be compared.

**Results:** We generated genome-wide ATAC-seq profiles of *A. calliptera* embryos across three developmental stages (3, 7 and 12 days post-fertilisation, dpf), identifying 550,876– 701,889 accessible chromatin peaks per stage. Accessible chromatin was enriched at gene promoters and at both highly conserved and accelerated conserved noncoding elements at every stage. The functional repertoire of accessible-promoter genes shifted across development, from early patterning at 3 dpf to the broadest regulatory repertoire at 7 dpf, and an increasingly neural- and locomotory-behaviour-enriched repertoire by 12 dpf, coinciding with post-hatching motor maturation. Differential accessibility showed that chromatin opening, rather than closing, dominates the earliest transitions (3 vs 7 and 3 vs 12 dpf), driven mostly by general metabolic genes, whereas the smaller, later 7 vs 12 dpf transition combined targeted gains at tissue-remodelling and apoptosis genes with losses at genes governing early neural and retinal patterning; almost all of these losses (96%) were at promoters that had opened between 3 and 7 dpf, indicating transient rather than sustained opening. Transcription factor (TF) footprinting identified stage-specific regulatory activity among developmentally important TFs, while the broader TF-promoter network remained topologically stable across all three stages, built around conserved hub regulators of core neural and sensory patterning genes e.g., SOX2, FOXG1, PTF1A, SOX10. This suggests that wiring is established early and largely maintained, following the hourglass model, with developmental progression driven by changes in activity of this conserved programme.

**Conclusions:** Together, these findings establish the first embryonic epigenomic resource for *A. calliptera*, revealing a developmental chromatin landscape of broad early opening and progressive functional specialisation, and providing a comparative framework for understanding how regulatory evolution has shaped Lake Malawi cichlid diversification.

## Introduction

East African cichlid fishes represent one of the most striking examples of an adaptive radiation, with rapid speciation and extensive phenotypic diversification across the African Great Lakes, particularly Lake Malawi [1]. Within this system, hundreds of species have evolved over a short evolutionary timescale (< 1 million years) and display striking variation in morphology, behaviour, and ecological niche specialisation [1–3]. Although this diversity has been linked to genomic variation, introgression, and ecological selection [3–5], the regulatory mechanisms that translate genetic variation into phenotypic divergence remain less well understood. In particular, early developmental regulatory landscapes, which could establish the trajectories from which adaptive traits emerge [6, 7], have received comparatively limited attention in Malawi cichlids.

Addressing this gap requires a system that captures regulatory features relevant to the ancestral state of the Lake Malawi radiation [8]. *Astatotilapia calliptera*, a riverine generalist inhabiting diverse habitats within and around Lake Malawi, is ideal because it is phylogenetically nested within the radiation yet retains shared ancestry with multiple derived lineages [2]. Its ecological flexibility and evolutionary position suggest that it may preserve regulatory programmes [9] central to the developmental potential from which more specialised cichlid lineages diversified. This makes *A. calliptera* a valuable system for investigating how developmental gene regulation may contribute to the origins of adaptive radiation [8, 9].

Genome-scale epigenomic approaches, particularly ATAC-seq, provide a powerful means to study how *cis-*regulatory architecture shapes vertebrate development [6, 10]. By identifying accessible chromatin regions, including promoters, and embedded transcription factor binding sites (TFBSs) [11], ATAC-seq can characterise stage-specific regulatory programmes that direct lineage commitment and tissue differentiation [10, 12, 13]. In teleosts [10] and cichlids[6, 14], chromatin accessibility studies have begun to link regulatory variation with trait diversification, including work in tilapia gill tissue [14] and broader comparative cichlid epigenomic analyses [6], but these and other studies [15] have focused on adult tissues or limited developmental contexts rather than a developmental time course in an ancestral-like Malawi lineage. A temporally resolved map of chromatin accessibility during early *A. calliptera* embryogenesis is therefore still lacking. Such a map is important because early embryogenesis is the period in which gene regulatory programmes first establish developmental trajectories, potentially influencing later phenotypic plasticity and adaptive capacity [10, 16]. Defining this landscape in *A. calliptera* would not only provide a foundational developmental epigenomic resource for cichlids, but also help identify regulatory features that may have broader relevance to the diversification of the Malawi radiation.

In this study, we generated a genome-wide map of chromatin accessibility across three embryonic stages of *A. calliptera* development, at 3, 7, and 12 days post-fertilisation (dpf), using ATAC-seq. These time points correspond to the pharyngula period (3 dpf; stages 14–16), the transition from embryonic to prejuvenile development shortly after hatching (7 dpf; approximately stage 20, when the oral jaws become functional) and a later prejuvenile stage (12 dpf; approximately stage 24) of the *A. calliptera* staging series [17]. They therefore bracket hatching (∼5 dpf), after which differences in head and jaw morphology, body shape and pigmentation between Malawi cichlid species become increasingly apparent, and precede yolk absorption and the onset of active feeding (∼16 dpf) [17].

Accessible regions were characterised according to their genomic context, with particular emphasis on non-coding regulatory elements and promoter-associated peaks, and stage-specific enrichment analyses were used to identify developmental programmes associated with each stage. Differential accessibility analyses defined regulatory changes across developmental transitions, and transcription factor footprinting was used to infer stage-dependent activity of t*rans*-acting regulators. We found that accessible chromatin is already pervasive at the earliest stage sampled and enriched at both highly conserved and accelerated conserved noncoding elements. The functional repertoire of accessible promoters is broadest at 7 dpf, shortly after hatching, and shifts towards neural and locomotory functions by 12 dpf. Notably, promoter opening, rather than closing, dominates the transition from 3 to 7 dpf, with transient opening at neural and retinal genes (e.g. *neurod4*, *irx7*, *notch1b* and *isl2b*), whereas the TF–promoter network is stable across the three stages: 822 of the 1,442 TF motifs tested (99.8% of those enriched) were significantly enriched at all three stages, none were stage-specific, and the same hub regulators, including SOX2, FOXG1, PTF1A and SOX10, were retained throughout. Together, these analyses establish a developmental epigenomic framework for *A. calliptera* and provide insight into regulatory mechanisms that may contribute to the adaptive potential of a generalist lineage associated with rapid cichlid diversification.

## Materials And Methods

### Tissue dissection and embryo staging

All animal procedures were approved by the University of Hull, UK, and carried out in accordance with approved guidelines. *Astatotilapia calliptera* ‘Ruvuma’ embryos were collected from the ovaries adult females and at different stages (3 dpf, 7 dpf, and 12 dpf), suspended in pre-chilled 1× PBS prior to dissection. For 3 dpf embryos, the animal pole was capped. For 7 dpf and 12 dpf embryos, the yolk sac was removed, and whole embryo taken for nuclei preparation. Embryos were staged based on collection date and developmental status.

### Nuclei preparation

Approximately 50,000 cells were pooled for each sample. Nuclei were pelleted at 500×g for 5 min at 4°C, washed once with ice-cold 1× PBS, and resuspended in cold lysis buffer (10 mM Tris-HCl pH 7.4, 10 mM NaCl, 3 mM MgCl2, 0.05% IGEPAL CA-630). Following lysis, nuclei were immediately pelleted and kept on ice until transposition.

### Transposition and library preparation

Pelleted nuclei were resuspended in transposition mix containing 2× TD buffer (Illumina), Tn5 transposase (Illumina), and nuclease-free water, and incubated at 37°C for 30 min. Transposed DNA was purified using the MinElute PCR purification kit (QIAGEN) and eluted in 10 mM Tris-HCl, pH 8. ATAC libraries were amplified using NEBNext High-Fidelity 2× PCR Master Mix (NEB) with indexed P5 and P7 adapters, then purified with MinElute and an additional AMPure XP clean-up step as described previously [6, 14]. Libraries were quantified using a Qubit 4 fluorometer and size distribution was assessed using an Agilent Tapestation and/or Bioanalyser.

### Naked DNA controls

Genomic DNA was extracted from pooled tissue using the DNeasy Blood and Tissue kit (QIAGEN) and used to prepare naked DNA libraries with the Nextera XT DNA Library Preparation kit (Illumina) according to the manufacturer’s protocol. These libraries were used as controls to estimate background accessibility and Tn5 sequence bias.

### Sequencing

ATAC-seq and naked DNA libraries were pooled equimolarly and sequenced as 50 bp paired-end reads on an Illumina NovaSeq 6000 platform at the Earlham Institute.

### ATAC-seq processing

Raw reads were trimmed with Trim Galore! (v0.5.0; https://github.com/FelixKrueger/TrimGalore) and quality checked using FastQC (v0.11.9) [18] with default settings. Reads were aligned to the *Astatotilapia calliptera* reference genome (fAstCal1.2) using Bowtie2 (v2.2.6) [19] with parameters --k 4 -- X2000 --mm, and output in BAM format with SAMtools (v1.9) [20]. Mitochondrial reads were identified by BLAST (v2.3.0) [21] and removed using SAMtools (v1.9) [20]. Duplicates were marked with Sambamba v0.6.5 [22], and low-quality, unmapped, non-primary, and failing-QC reads were filtered using SAMtools (v1.9) [20], retaining properly paired alignments. BAM files were converted to tagAlign format using Bedtools (v2.30.0) [23], Tn5-shifted, and peaks were called with MACS2 (v2.1.1) [24, 25] using the corresponding naked DNA control as input. Coverage tracks were generated using UCSC tools (v333; http://hgdownload.cse.ucsc.edu/admin/exe/), and replicate concordance was assessed using the Irreproducible Discovery Rate (IDR). Consensus peak sets were defined from reproducible peaks across biological replicates, following the ENCODE ATAC-seq pipeline (https://www.encodeproject.org/atac-seq/). Fragment length distributions were visualised with Picard (v1.140; https://github.com/broadinstitute/picard/), and TSS enrichment was calculated with ATACseqQC (v1.18.0) [26]. Only samples passing standard ATAC-seq quality thresholds were retained for downstream analyses.

### Peak annotation and genomic feature analysis

Consensus peaks from both biological replicates of each stage were intersected with the *A. calliptera* (fAstCal1.2) genome annotation using Bedtools (v2.30.0) [23] to assign them to promoters, introns, exons, intergenic regions, UTRs and conserved non-coding elements (hCNEs and aCNEs; see below). A peak overlapping more than one feature was counted for each. Promoters were defined as the 5 kb upstream of the transcription start site (TSS) on the gene’s strand, shortened where a neighbouring gene lay closer than 5 kb. The core promoter was taken as ±100 bp around the TSS. To assess promoter-proximal accessibility, each peak summit was classified by its distance to the nearest TSS (≤100 bp, 100 bp–5 kb, or >5 kb). For each class, peak density was calculated as the number of peaks per kb of search space, defined as the total length of the assembly lying within the class distance of any annotated TSS, and the number of genes per class was obtained by assigning each peak summit to its nearest TSS. Enrichment of peaks in each feature was tested by permutation: peaks, merged and restricted to the numbered linkage groups, were randomly repositioned within their chromosomes 200 times (bedtools shuffle) to give the expected overlap. Fold enrichment is the observed over the mean expected count, and empirical p-values were Benjamini-Hochberg (BH) corrected [27], with q < 0.05 considered significant.

### Functional enrichment analysis

Gene Ontology (GO) enrichment of biological processes (GO:BP) was carried out with the g:GOst module of g:Profiler [28], using the *A. calliptera* annotation, GO:BP as the only data source, all annotated genes as the statistical background, and the Benjamini-Hochberg (BH) false discovery rate (FDR) correction; terms with an adjusted p < 0.05 were considered significant. For each developmental stage, the query was the set of genes whose 5 kb promoter overlapped an ATAC-seq peak (gprofiler2 v0.2.4). For differential accessibility (below), the queries were the genes gaining or losing promoter accessibility in each pairwise comparison (gprofiler2 v0.2.4). Fold enrichment was calculated as (query genes in the term / query size) / (term size / background size).

### Differential chromatin accessibility

Differential accessibility between developmental stages was assessed using csaw (v1.42.0) [29] and edgeR (v4.6.3) [30]. Read counts were summarised in 150 bp sliding windows (50 bp spacing) across primary linkage groups for each pairwise stage comparison, retaining windows with signal >3-fold above a 2 kb local background estimate, and normalised for composition bias using background bin counts. Differential testing used an edgeR quasi-likelihood F-test (BH FDR < 0.05); adjacent significant windows within 100 bp were merged into consolidated differentially accessible regions using csaw::mergeWindows and combineTests (region-level Simes-combined FDR < 0.05). Differentially accessible regions were then overlapped with promoter annotations to define gene-associated changes. Overlaps among the sets of genes with accessible promoters at each stage, and between these sets and differentially accessible genes, were computed from gene identifiers and visualised as a Venn diagram. Genes linked to stage-specific promoter accessibility were subsequently subjected to GO enrichment analysis as above.

### Multiple genome alignment and identification of conserved noncoding elements (CNEs)

A reference-free multiple genome alignment (MGA) of *A. calliptera* (fAstCal1.2) and five cichlid genomes - *M. zebra* UMD2a [31, 32]; *P. nyererei* v1 [33]; *A. burtoni* v1 [33]; *N. brichardi* v1 [33]; *O. niloticus* UMD_NMBU [32, 34] *-* was created using Cactus (v2.0.3) [35] and outputted in Multiple Alignment Format (MAF). To identify CNEs, a neutral substitution model was created using the *phyloFit* function of PHAST (v1.5) [36] by fitting a time reversible substitution ‘REV’ model and parameters ‘—tree “(((((Metriaclima_zebra, Astatotilapia_calliptera),Pundamilia_nyererei),Astatotilapia_burtoni),Neolamprologus_brichardi),Oreochromis_niloticus)” --subst-mod REV’. The six cichlid MGA was split by *O. niloticus* chromosome/scaffold using the *mafSplit* function in the UCSC-tools package (v333; http://hgdownload.cse.ucsc.edu/admin/exe/). The neutral substitution model and MGAs of each *O. niloticus* reference chromosome/scaffold were used as input to predict conserved noncoding elements (CNEs) using the *phastCons* function of PHAST (v1.5) [36] with parameters ‘--target-coverage 0.3 --expected-length 30 -- most-conserved --estimate-trees --msa-format MAF’. The conservation score output was used to define highly conserved CNEs (hCNEs) with sequence identity ≥90% over ≥30bp and pseudo accelerate/diverged CNEs (pseudo aCNEs) with sequence identity <90% over ≥30bp. The evolutionary conservation-acceleration (CONACC) score of each pseudo aCNE was predicted by running *phyloP* [37] of the PHAST (v1.5) package [36] using the likelihood ratio test (LRT) ‘--method LRT’ on the CNE -- features in ‘--mode CONACC’ with the neutral substitution model of the corresponding pairwise *O. niloticus–A. calliptera* alignment, such that the test was run separately for each species against *O. niloticus*. Pseudo aCNEs with a negative CONACC score, likelihood ratio <0.05, and significant divergence (altsubscale >1), were defined as significantly deviating from the neutral model, and therefore as true aCNEs. The test therefore identifies elements that have diverged from neutral expectation between *O. niloticus* and *A. calliptera* and does not assign the acceleration to a particular branch. Since CNEs were defined by their position in *O. niloticus* chromosomes as a reference, corresponding CNE coordinates in *A. calliptera* scaffolds were defined using a combination of the *mafsinRegion* function of the UCSC-tools package (v333; http://hgdownload.cse.ucsc.edu/admin/exe/) and Bedtools (v2.30.0) [23].

### Transcription factor footprinting and motif analysis

TF footprints were identified in the pooled replicate ATAC-seq data of each stage using HINT-ATAC in the Regulatory Genomics Toolbox, and motif matching within footprints used a stringent false positive rate (FPR) of 0.0001. The position weight matrices (PWMs) were species-specific and cichlid-wide matrices defined in our previous study [38], together with vertebrate PWMs from JASPAR v9.0 [39, 40] and HOCOMOCO [41]. Enrichment of motif-predicted binding sites in genomic features was tested with the Genomic Association Tester (GAT) [42], using the footprint-supported motif matches of each stage as segments, the genomic feature annotation (promoters, exons, introns, UTRs, intergenic regions, hCNEs and aCNEs) as annotations, chromosome lengths as the workspace, and BH correction [27] (q < 0.05). TF activities were summarised using the mean motif-match bit-score across gene promoters. K-means clustering was explored for k=1 to 20, and the optimal number of clusters was selected using the elbow criterion and silhouette width, with average silhouette values above 0.25 considered indicative of structured clustering. Bit-scores were *Z-score* transformed by row and visualised as a heatmap in R (v4.5.2) using the ComplexHeatmap package (v3.6). TF footprint plots were generated from bias-corrected signals using the differential analysis module of HINT-ATAC in the Regulatory Genomic Toolbox (v0.13.0) package [43].

### TF–promoter regulatory network topology

To characterise the topology of the TF–promoter regulatory network inferred from footprint-supported motif enrichment, each TF motif significantly enriched (BH FDR < 0.05) in accessible gene promoters at a given stage was treated as a network hub node, with directed edges to every target gene promoter reported for that motif by enrichAnalyzer (https://github.com/Roy-lab/enrichAnalyzer_Nongraph). For each stage the following was recorded: the number of significantly enriched TF motifs (network breadth); the promoter-target (‘out-degree’) count and fold enrichment for each motif, used to rank hub regulators; and, using set intersection on motif identity across the three stages, the number of motifs shared across all stages versus stage-restricted motifs. This analysis is distinct from the TF–target gene examples described above, which draw on TF footprints located within accelerated conserved noncoding elements (aCNEs) specifically; the network topology analysis instead considers all significantly enriched promoter-associated motifs genome-wide, irrespective of CNE overlap.

## Results

### Accessible chromatin is pervasive throughout *A. calliptera* embryogenesis and enriched at conserved regulatory elements

To characterise the regulatory landscape during early embryogenesis in *A. calliptera*, we performed high-depth ATAC-seq of embryos at three key developmental stages (3, 7, and 12 days post-fertilisation, dpf; two biological replicates per stage; Fig. 1A), identifying 701,889, 550,876 and 672,569 open chromatin (accessible) peaks at 3, 7 and 12 dpf, respectively (union of peaks called in either biological replicate at each stage; see *‘Materials and Methods’*; Fig. 1B; Supplementary Table S1a). These accessible peaks could be associated with 24,156, 21,550, and 23,474 of the 24,198 annotated genes (99.8%, 89.1% and 97.0%) at 3, 7 and 12 dpf, respectively. We first annotated (Supplementary Table S1) and tested the enrichment of accessible peaks across coding and noncoding genomic regions in the *A. calliptera* fAstCal1.2 reference genome (see ‘*Materials and Methods*’). Accessible peaks were significantly enriched (FDR < 0.05) across multiple genomic features, with enrichment patterns varying between developmental stages (Fig. 1C). The highest enrichments were observed for 5′ UTR regions at 7 dpf (fold enrichment = 3.31) and 12 dpf (fold enrichment = 2.38), while highly conserved noncoding elements (hCNEs, see ‘*Materials and Methods*’) were enriched across all stages (3 dpf: fold enrichment = 1.84; 7 dpf: fold enrichment = 2.32; 12 dpf: fold enrichment = 1.96). Additional enrichment was observed for accelerated conserved noncoding elements (aCNEs, see ‘*Materials and Methods*’) at 3 dpf (fold enrichment = 1.60), 7 dpf (fold enrichment = 1.53), and 12 dpf (fold enrichment = 1.67), and coding exons at all stages (3 dpf: fold enrichment = 1.59; 7 dpf: fold enrichment = 1.66; 12 dpf: fold enrichment = 1.63) (Fig. 1C).

**Figure 1.**
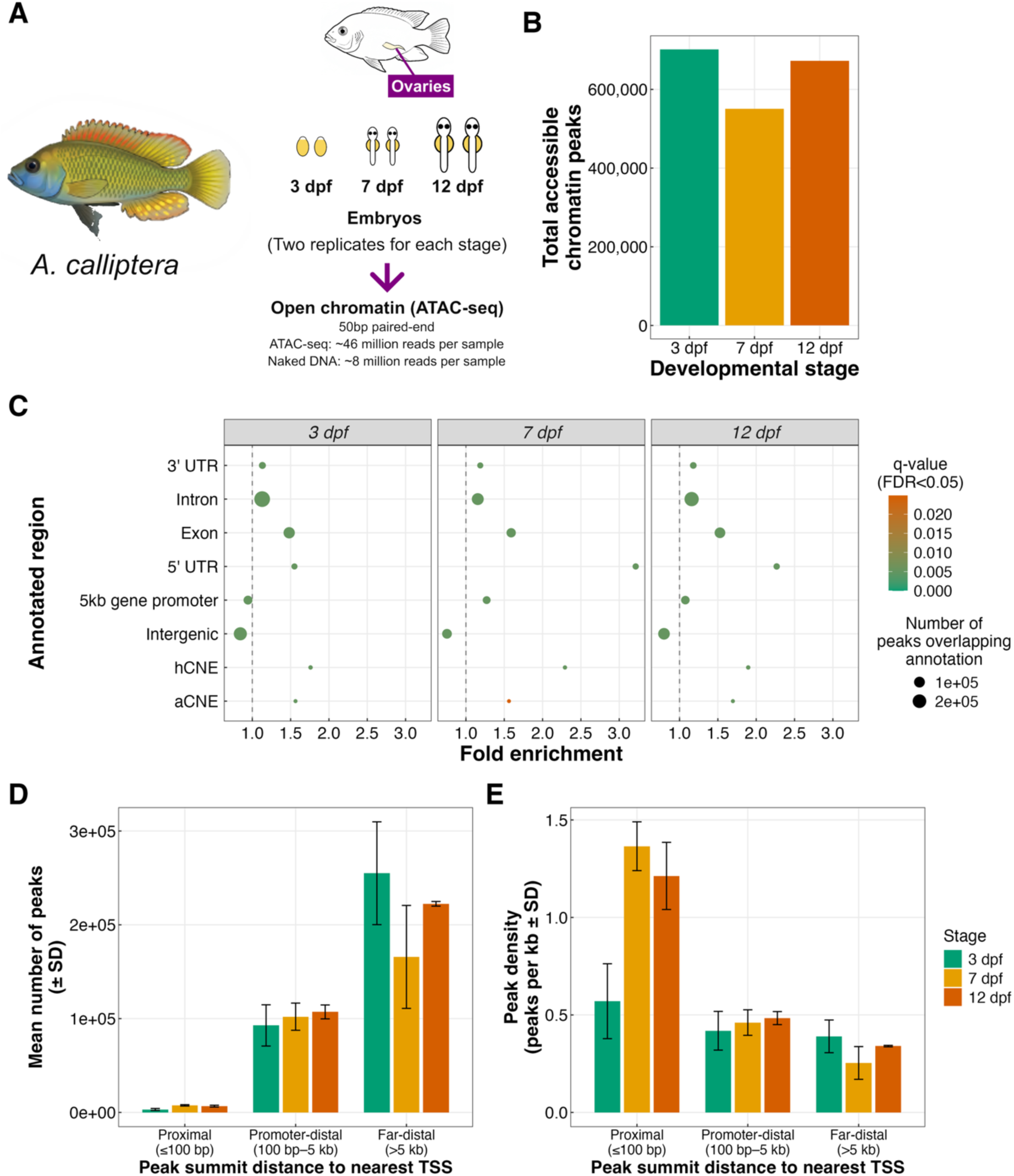
Chromatin accessibility landscape across *A. calliptera* embryonic development. **(A)** Experimental design schematic: whole embryos were collected from two biological replicates at each of three developmental stages (3, 7 and 12 days post-fertilisation, dpf) for ATAC-seq (50-bp paired-end reads; ∼46 million reads per sample), alongside naked genomic DNA background controls (∼8 million reads per sample). **(B)** Total number of accessible chromatin peaks identified at each developmental stage (union of peaks called in either biological replicate; *see ‘Materials and Methods’*). **(C)** Enrichment of accessible peaks across annotated genomic features at each stage, assessed by permutation testing (200 genome-wide segment shuffles per stage); circle size indicates the number of peaks overlapping each annotation and colour indicates the Benjamini-Hochberg-corrected q-value (FDR<0.05). **(D)** Mean number of accessible peaks per stage by summit distance to the nearest TSS, split into promoter-proximal (≤100 bp), promoter-distal (100 bp–5 kb) and far-distal (>5 kb) categories (error bars: SD across the two biological replicates; Supplementary Table S1). **(E)** Peak density, i.e. the counts in (D) divided by the search space of each class (the length of the assembly lying within that distance of any annotated TSS: 5.5 Mb, 221.5 Mb and 653.4 Mb for proximal, promoter-distal and far-distal regions, respectively), expressed as peaks per kb (error bars: SD across the two biological replicates; Supplementary Table S1E).

To link accessible peaks to gene functions, we next examined their distribution relative to the nearest gene transcription start site (TSS), distinguishing promoter-proximal peaks (≤100 bp from the TSS), promoter-distal peaks (100 bp–5 kb from the TSS, i.e. within a 5 kb promoter region), and far-distal peaks (>5 kb from the TSS) (Fig. 1D; Supplementary Table S1b-d) as done previously [6, 38]. Far-distal peaks were the highest numerically at every stage, with mean counts of 255,000 ± 54,795 at 3 dpf, 165,795 ± 54,864 at 7 dpf, and 222,392 ± 2,481 at 12 dpf. Promoter-distal peaks (100 bp–5 kb from the TSS) were comparable across stages (92,782 ± 21,975 at 3 dpf; 102,074 ± 14,492 at 7 dpf; 107,168 ± 7,398 at 12 dpf), while promoter-proximal peaks (≤100 bp) were the least numerous at every stage (3,163 ± 1,065 at 3 dpf; 7,570 ± 694 at 7 dpf; 6,725 ± 955 at 12 dpf) (Fig. 1D; Supplementary Table S1b-d). These counts reflect the size of each search space: far-distal regions comprise 74.2% of the assembly, promoter-distal regions 25.2% and promoter-proximal regions only 0.6%. Normalised for search space, promoter-proximal peaks were the most densely distributed class at 7 and 12 dpf (1.37 and 1.21 peaks per kb, versus 0.46 and 0.48 in promoter-distal and 0.25 and 0.34 in far-distal regions), whereas at 3 dpf peak density was more similar across classes (0.57, 0.42 and 0.39 peaks per kb, respectively; Fig. 1E; Supplementary Table S1E). The number of genes associated with each class likewise does not simply track peak number: assigning each peak summit to its nearest gene TSS, promoter-proximal peaks mapped to 4,975, 7,996 and 7,853 genes at 3, 7 and 12 dpf (20.6%, 33.0% and 32.5% of the 24,198 annotated genes), compared with 21,389, 18,869 and 20,665 genes (88.4%, 78.0% and 85.4%) for promoter-distal peaks, as the promoter-proximal window spans only 200 bp around the TSS (Supplementary Table S1F). These patterns indicate that, although most peaks lie far from TSSs, reflecting the size of that search space, peak density is highest at promoter-proximal regions at 7 and 12 dpf (Fig. 1E) and a substantial fraction of peaks fall within the broader 5 kb promoter region, and likely harbour gene regulatory motifs as identified in cichlids previously [6, 14, 38, 44].

We focused on gene promoter regions, defined as regions within 5 kb upstream of a genes TSS, as these regions harbour regulatory elements including transcription factor binding sites (TFBSs) that can influence developmental gene regulation. Across the three developmental stages, promoter-associated accessible peaks were identified for 67,563 peaks associated with 17,585 genes at 3 dpf, 114,240 peaks associated with 16,101 genes at 7 dpf, and 101,969 peaks associated with 18,329 genes at 12 dpf (Supplementary Table S1b). Most of these genes were shared between stages: of the 20,418 genes with an accessible promoter in at least one stage, 13,540 (66.3%) had one at all three stages, whereas 1,580, 201 and 580 genes were unique to 3, 7 and 12 dpf, respectively, and 308, 2,157 and 2,052 were shared by only 3 and 7, 3 and 12, and 7 and 12 dpf, respectively (Supplementary Fig. 2A; Supplementary Table S1G). As a specificity control, we also tested genes with no promoter-overlapping accessible peak at each stage (6,613 genes at 3 dpf, 8,097 at 7 dpf, and 5,869 at 12 dpf) for gene ontology (GO) enrichment against the same background (Supplementary Table S1c): at every stage these genes were strongly and specifically enriched for immune system process and immune response (adj. p < 10⁻²⁰ at every stage), rather than any developmental term (Supplementary Figure 1). These genes are potentially inactive prior to the development of an adaptive immune system [45], and support the specificity of the developmental enrichment reported below for genes with accessible promoters. These promoter-associated accessible regions therefore represent a largely shared promoter regulatory landscape with a smaller stage-specific component, with the highest number of promoter peaks detected at 7 dpf and the greatest number of associated genes detected at 12 dpf.

### Stage-specific enrichment of regulatory elements and functional annotations

GO enrichment analysis of genes associated with promoter-overlapping peaks identified core shared significant terms present at every developmental stage (Fig. 2A; Supplementary Table S2), including developmental process (fold enrichment = 1.09– 1.13, adj. p = 1.77×10⁻⁴² at 3 dpf, 3.79×10⁻⁵⁹ at 7 dpf, 3.08×10⁻⁵⁰ at 12 dpf), anatomical structure development (fold enrichment = 1.08–1.13), multicellular organism development, cell differentiation, nervous system development and neurogenesis. Sensory-system and eye-related terms were also significant at every stage, including sensory system development (adj. p = 7.2×10⁻⁴, 4.4×10⁻⁸ and 4.5×10⁻⁶ at 3, 7 and 12 dpf) and retina development in camera-type eye (9.0×10⁻⁴, 2.7×10⁻⁶, 2.4×10⁻⁴) (fold enrichment = 1.11–1.17; Supplementary Table S2). Genes annotated to the latter include *dnmt1*, required for retinal stem cell and lens maintenance in zebrafish [46, 47], and *otx2a*, an ortholog of *otx2*, which is required for optic vesicle and retinal pigment epithelium development [48] and whose expression in the fore- and midbrain and eyes and differential regulation among Malawi lineages has been documented by *in situ* hybridisation [7]. Enrichment of nervous-system-related terms across all three stages is consistent with the continuous expression of neurogenesis, forebrain-patterning and sensory-system genes reported across equivalent pharyngula and larval stages of Lake Malawi cichlid development [7], and more broadly with chromatin accessibility supporting a persistent core developmental and neural programme throughout early vertebrate embryogenesis [16, 49].

**Figure 2.**
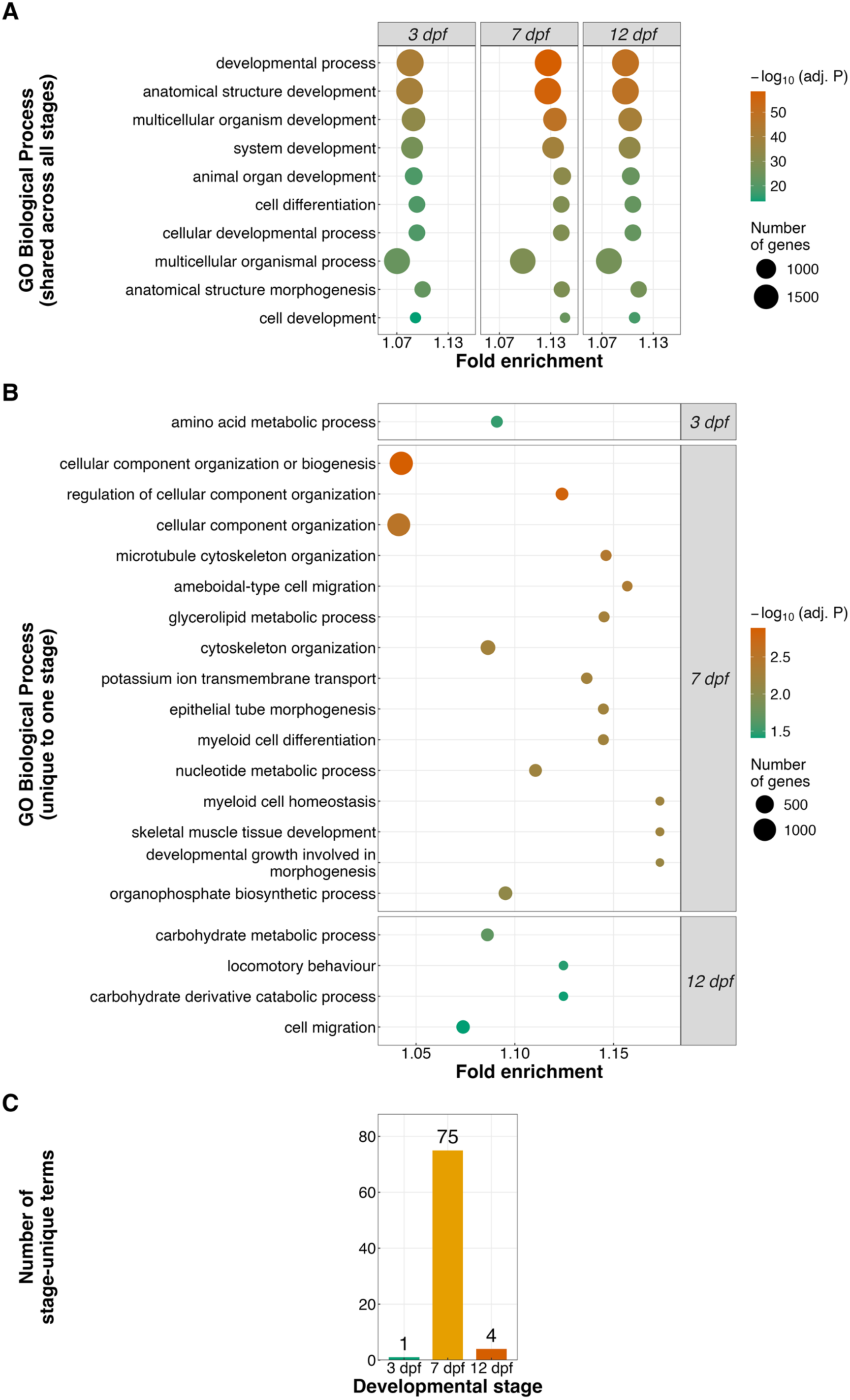
Gene Ontology (GO) Biological Process enrichment of genes with accessible gene-promoter peaks across developmental stages. (A) The ten most significant GO:BP terms shared across all three stages (ranked by smallest adj. p-value across stages), shown for 3, 7 and 12 dpf. (B) The 15 most significant GO:BP terms unique to a single stage (all terms where fewer than 15 were unique: n = 1 at 3 dpf and n = 4 at 12 dpf). In (A) and (B), the x-axis shows fold enrichment, circle size shows the number of query genes annotated to the term, and colour shows −log₁₀ adj. p (g:Profiler g:GOst, adj. p < 0.05; background = all annotated *A. calliptera* genes). **(C)** Number of GO:BP terms significantly enriched at only one developmental stage. Full results are in Supplementary Table S2.

Beyond this shared core, the set of stage-restricted GO:BP terms differed markedly in both number and content across development (Fig. 2B-C; Supplementary Table S2). As most promoter-associated genes are shared between stages (13,540 of 20,418; Supplementary Fig. 2A), these stage-restricted terms are likely to reflect differences in enrichment among largely overlapping gene sets rather than stage-exclusive gene sets; for example, only 201 genes were unique to 7 dpf, compared with 75 stage-unique terms. The 3 dpf stage contained a single stage-unique term, amino acid metabolic process (fold enrichment = 1.09, adj. p = 0.032), indicating a core metabolic component at this earliest stage (Fig. 2B-C; Supplementary Table S2). The 7 dpf stage showed the greatest number of stage-unique terms (n = 75), consistent with 7 dpf capturing the broadest regulatory programme of the three stages; dominated by organogenesis processes, including neural retina development (adj. p = 0.017) and photoreceptor cell differentiation (adj. p = 0.026), hindbrain development (adj. p = 0.010), inner ear development (adj. p = 0.038), and somite development (adj. p = 0.017) (Fig. 2B-C; Supplementary Table S2), consistent with the onset of sensory organ, craniofacial and musculoskeletal differentiation reported for Malawi cichlid embryos around this stage [9]. The 7 dpf embryos approximately correspond to stage 20, shortly after hatching (∼5 dpf), when the oral jaws first become functional [17]. A further, fish-specific example among the 7 dpf-unique terms was lateral line development and neuromast development (adj. p = 0.045 and 0.033) (Fig. 2B-C; Supplementary Table S2), reflecting active patterning of the mechanosensory lateral line system at this stage — a timeline consistent with the known emergence and distribution of superficial neuromasts in larval cichlids [50]. By 12 dpf, the stage-unique term set was again small (n = 4) but distinct, comprising locomotory behaviour (fold enrichment = 1.12, adj. p = 0.035), cell migration (adj. p = 0.039), and carbohydrate metabolism (carbohydrate metabolic process and carbohydrate derivative catabolic process, adj. p = 0.021 and 0.038) (Fig. 2B-C; Supplementary Table S2). This is consistent with post-hatching locomotor and metabolic maturation at approximately stage 24 (11–13 dpf), by which time *A. calliptera* embryos swim upright but have not yet fully absorbed the yolk or begun active exogenous feeding (∼16 dpf) [17], and with the general transition from yolk-based to exogenous nutrition in teleost larvae [51].

Together, these results indicate that a conserved core developmental and neurogenic programme is maintained throughout early *A. calliptera* embryogenesis, with the specific functional repertoire diversifying substantially by 7 dpf into distinct sensory, craniofacial and musculoskeletal organogenesis processes, and narrowing again by 12 dpf towards behavioural and metabolic maturation, broadly consistent with the conserved trajectory of vertebrate embryonic development [7, 9]

### Differential chromatin accessibility between developmental stages

To identify regulatory elements associated with developmental transitions in *A. calliptera*, differential chromatin accessibility (FDR < 0.05) was assessed between embryonic stages, identifying stage-specific chromatin remodelling across development. The largest number of differentially accessible regions was observed between 3 and 7 dpf (26,102 regions), followed by 3 vs 12 dpf (21,759 regions), with fewer regions identified between 7 and 12 dpf (3,450 regions) (Fig. 3A; Supplementary Table S3), indicating that the greatest chromatin changes occur during early embryogenesis.

**Figure 3.**
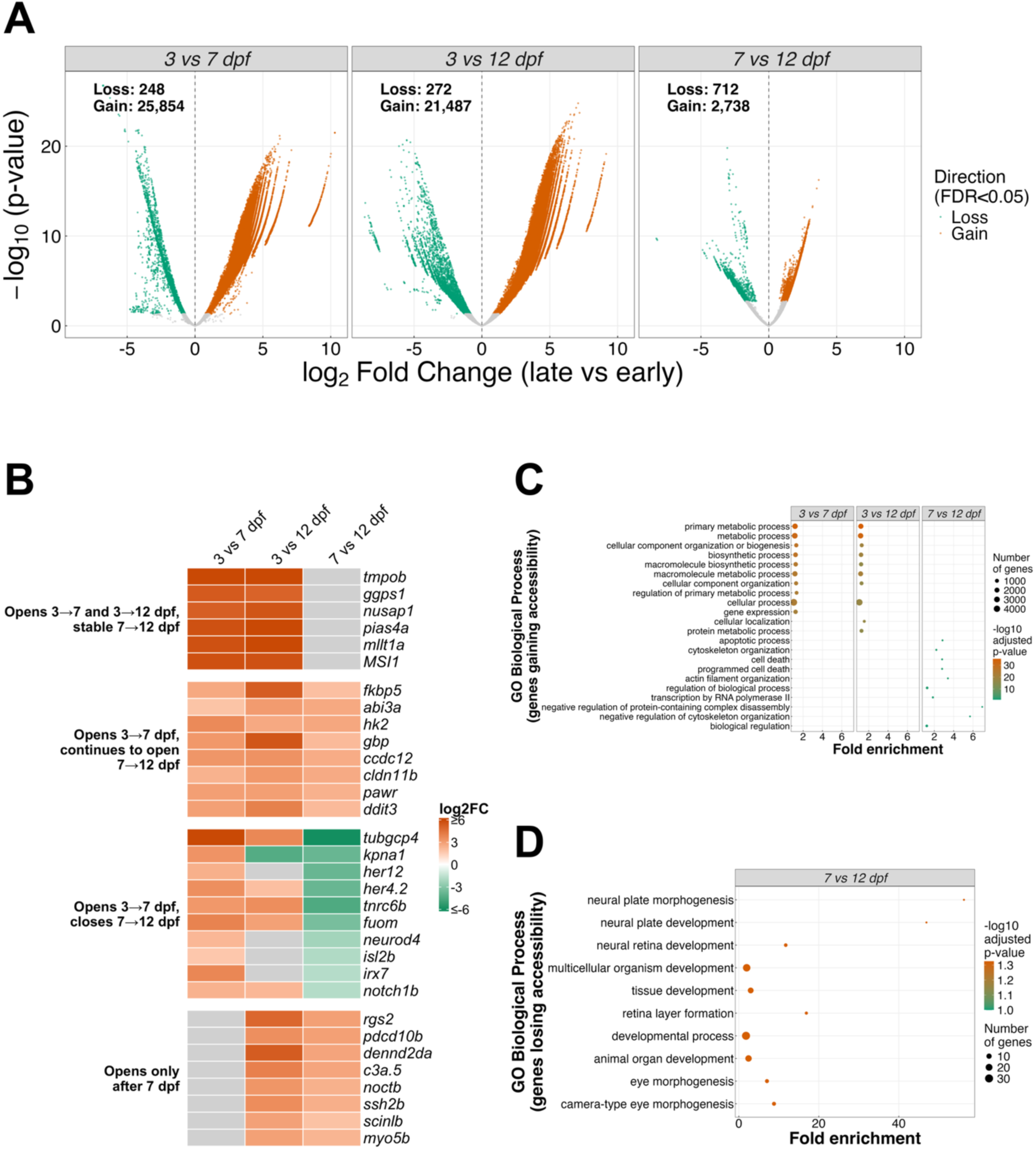
Differential chromatin accessibility between developmental stages. **(A)** Volcano plots of window-level differential accessibility for each pairwise stage comparison (log2 fold change vs -log10 p-value); points are coloured by significant direction of change (FDR<0.05). **(B)** Mean log2 fold change (later vs earlier stage) in accessibility of differentially accessible regions (FDR < 0.05) overlapping gene promoters, for annotated genes representing four temporal patterns: opening at 3 vs 7 and 3 vs 12 dpf with no later change; opening at 3 vs 7 dpf and continuing to open at 7 vs 12 dpf; opening at 3 vs 7 dpf then closing at 7 vs 12 dpf; and opening only after 7 dpf. The six most significant genes per pattern are shown (ranked by region-level FDR), plus genes discussed in the text (*pawr, ddit3, neurod4, irx7, notch1b, isl2b, scinlb, myo5b*). Grey: no significant region overlapping the promoter in that comparison; colour scale capped at ±6. **(C)** GO Biological Process enrichment of genes gaining promoter accessibility, and **(D)** genes losing promoter accessibility, shown for up to the top 10 terms per comparison.

Given the relationship between promoter accessibility and transcriptional regulation, downstream analyses focused on regions overlapping ±5 kb promoter regions, based on TFBS enrichment and activity in this genomic window in five closely-related cichlids [6, 38]. Promoter-associated differential accessibility followed the same overall pattern, with 9,606 differential regions in the 3 vs 7 dpf comparison, 8,380 regions in the 3 vs 12 dpf comparison, and 799 regions in the 7 vs 12 dpf comparison, mapping to 8,912, 7,701 and 793 unique genes, respectively (Supplementary Table S3A). Promoter accessibility changes were strongly biased towards gains: 8,878 genes gained and only 37 lost accessibility at 3 vs 7 dpf (99.6% gains; median log2 fold change 3.3), and 7,664 gained versus 40 lost at 3 vs 12 dpf (99.5%; median 3.6), whereas the smaller 7 vs 12 dpf comparison was bidirectional, with 604 genes gaining (76%; median 1.6) and 190 losing (24%; median −2.3) accessibility (Supplementary Table S3B). These gene numbers should be read against the 13,848, 15,697 and 15,592 genes with an accessible promoter at both stages of the respective comparison: 7,420 (53.6%), 6,506 (41.4%) and 770 (4.9%) of those genes, respectively, showed differential promoter accessibility, while the remaining 1,492, 1,195 and 23 differentially accessible genes (16.7%, 15.5% and 2.9%) had an accessible promoter at only one of the two stages (Supplementary Table S3F). Of the 13,540 genes with an accessible promoter at all three stages, 7,900 (58.3%) were differentially accessible in at least one comparison, and genes accessible only at 7 and 12 dpf were most often differentially accessible (1,458 of 2,052; 71.1%), consistent with promoter opening after 3 dpf (Supplementary Fig. 2B; Supplementary Table S1G).

Among the 7 vs 12 dpf changes, most late gains (93%; 560/604 gene promoters) were also higher than at 3 dpf. Almost all late losses (96%; 182/190) occurred at promoters that had opened between 3 and 7 dpf (Supplementary Table S3C), indicating transient rather than sustained opening (Fig. 3B). Transiently open promoters included *neurod4*, *irx7*, *notch1b* and *isl2b* (Fig. 3B), genes that act at successive steps of vertebrate neurogenesis and retinal differentiation, from Notch-mediated maintenance of progenitors (*notch1b*) [52] to proneural and retinal-fate specification (*neurod4*, *irx7*) [53, 54] and retinal ganglion cell identity (*isl2b*) [55]. Promoter opening between 3 and 7 dpf followed by closure by 12 dpf is therefore consistent with a window of neural and retinal specification that is largely complete by 12 dpf. Because the visual system is a major axis of regulatory divergence among Lake Malawi cichlids [6, 38, 44, 56], these loci are candidates for lineage-specific regulatory change associated with cichlid development. In contrast, *ssh2b*, *scinlb* and *myo5b*, which opened only after 7 dpf (Fig. 3B), encode regulators of actin dynamics and epithelial morphogenesis [57–59], consistent with the cytoskeletal remodelling and cell-death signature of later embryogenesis.

GO enrichment of genes gaining promoter accessibility in the earliest transitions (3 vs 7 dpf and 3 vs 12 dpf) identified broad enrichment for core cellular and metabolic processes, including primary metabolic process, biosynthetic process and cellular component organisation (adj. p < 10⁻²⁰ for all terms) (Fig. 3C), consistent with widespread activation of general transcriptional and translational machinery during this period of rapid growth and tissue differentiation [60]. By the 7 vs 12 dpf transition, genes gaining promoter accessibility were enriched for apoptotic process and programmed cell death (adj. p = 0.006) and for cytoskeleton and actin filament organisation (adj. p = 0.006 and 0.009; Fig. 3D), including the pro-apoptotic genes *pawr* and *ddit3* and the actin-associated genes *ssh2b*, *scinlb* and *myo5b* [57–59], consistent with the tissue remodelling events of later embryogenesis. In contrast, genes losing promoter accessibility showed a modest enrichment (adj. p = 0.047) for neural plate morphogenesis and development (e.g., *nkd1*, *neurod4* genes) and neural retina development (e.g., *irx7*, *tubgcp4*, *exoc5*, *wrap73* genes) (Fig. 3D). This suggests that regulatory programmes governing early neural patterning could become less accessible once this patterning is complete however, as this is based on only a few genes it requires further testing using more comparative stages. Nonetheless, this pattern is consistent with the expression of neurod-family and Iroquois-family transcription factors in the developing brain and eyes of Malawi cichlid embryos at equivalent stages [7].

Overall, these data indicate that chromatin accessibility change is greatest during early embryogenesis and is dominated by widespread gains in promoter accessibility at genes supporting general metabolic and biosynthetic activity, while the more modest, 7 vs 12 dpf transition captures a more targeted regulatory shift, combining accessibility gains of tissue-remodelling genes with losses at early neural-patterning genes as embryonic development proceeds.

### Transcription factor footprinting reveals stage-specific regulatory programmes

Transcription factors are key drivers of gene regulatory innovations, shaping gene regulatory networks associated with cichlid phenotypic diversity [14, 38, 44] and vertebrate embryonic development [61]. To profile TF-mediated regulation, we mapped a mean of 1,227,888 TF footprints across 3, 7, and 12 dpf *A. calliptera* embryos (Supplementary Table S4.1, see ‘*Materials and Methods*’). Of these, 89,625 to 144,843 footprints were located in gene promoters across the three stages, using 1,442 unique TF motifs from cichlid [38] and vertebrate [40, 41] lineages. These footprints yielded 897,521 to 1,393,388 non-redundant TFBSs (Supplementary Table S4.1), with a mean of 1.2 million non-redundant promoter TFBSs per stage; this is broadly consistent with the 1.4 million TFBSs identified in five adult cichlid species [6] and with the 3.6 million footprints mapped in the approximately threefold larger human genome [62].

Consistent with previous findings [6, 14, 38, 44], TFBSs were significantly enriched (adj. p < 0.05) in non-coding regulatory regions, including gene promoters and conserved non-coding elements (CNEs), across all three stages (Fig. 4A; Supplementary Table S4.2). Relative to gene promoters, TFBSs were more than two-fold enriched in highly conserved CNEs shared with other East African cichlids (hCNEs; sequence identity ≥90% over ≥30 bp; see ‘*Materials and Methods*’), indicating that many binding sites are maintained by selective constraint (Fig. 4A; Supplementary Table S4.2). In contrast, TFBSs were less enriched in accelerated CNEs (aCNEs; sequence identity <90% over ≥30 bp and significant deviation from a neutral model; see ‘*Materials and Methods*’), highlighting a subset of putative stage-specific regulatory sites (Fig. 4A; Supplementary Table S4.2). Only 11 or fewer TF– target gene interactions involving aCNEs were shared across all stage comparisons; for example, the PAX5–ddx54 interaction in the 7 vs 12 dpf comparison may be associated with midbrain–hindbrain boundary patterning [63] (Supplementary Table S4.3). By contrast, up to 138 aCNE-linked TF–target gene interactions were unique to individual comparisons, including FOXA1–otud7a in 12 dpf vs 3 dpf, which may relate to organogenesis [64], and ALX1–tmem88b in 3 dpf vs 7 dpf, which may reflect neural crest migration [65] (Supplementary Table S4.4).

**Figure 4.**
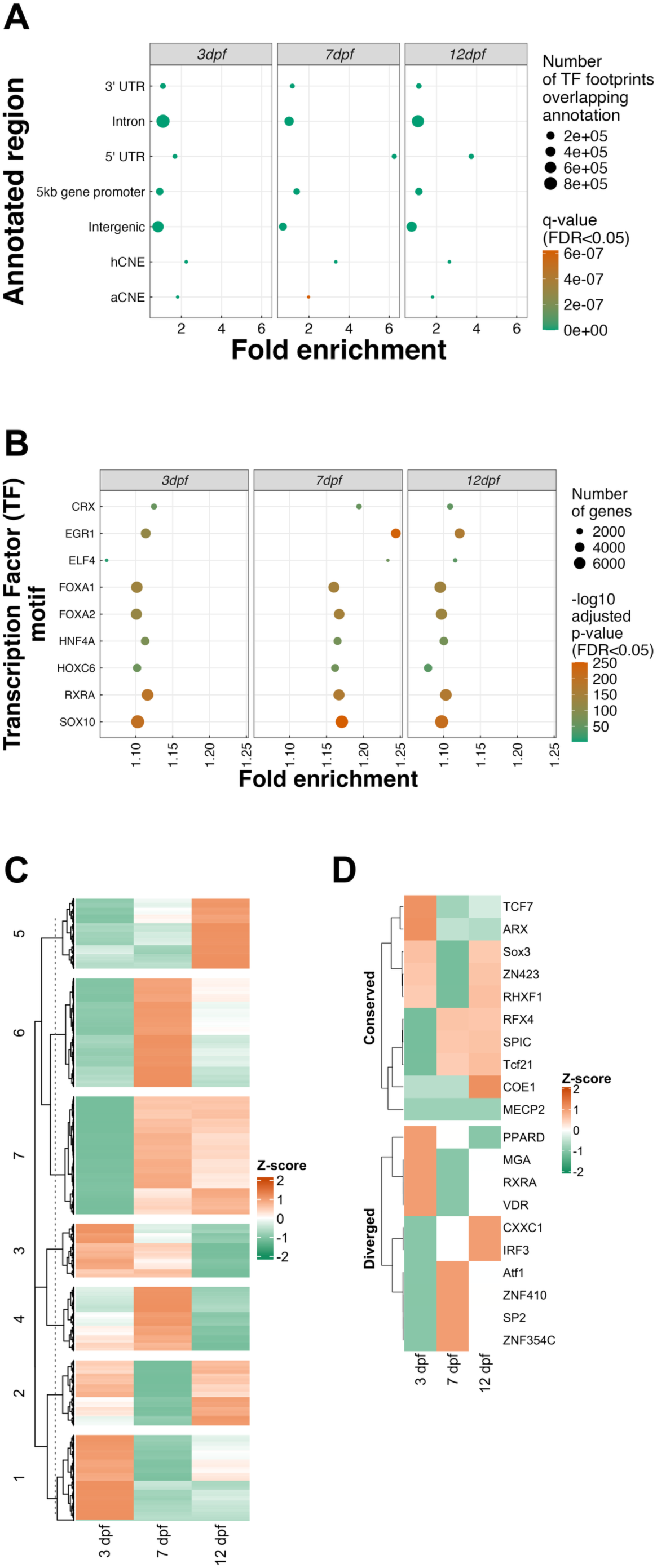
TF motif enrichment and activity landscapes in *A. calliptera* embryos. **(A)** Fold enrichment of TF motifs overlapping coding and noncoding regions across the three stages. Circles indicate enriched annotated regions at FDR < 0.05, with fold enrichment plotted on the x-axis and the number of overlapping TF motifs represented by circle size; **(B)** Enrichment of selected functionally associated TF motifs in gene promoter peaks across the three stages. Circles indicate TF motifs, with fold enrichment shown by the heatmap and significance plotted as -log10 FDR < 0.05 across species and tissues; circle size indicates the number of enriched genes per TF motif. **(C)** Heatmap of TF activities in gene promoter regions across developmental stages. Rows show clustered TFs and columns show species. Mean TF bit-scores across gene promoters, used as a proxy for activity, were Z-score transformed across rows; green indicates low activity and orange indicates high activity; **(D)** Heatmap of the top 10 most conserved and top 10 most diverged TF motifs in gene promoter regions across 3, 7 and 12 dpf. Mean motif bit-scores were Z-score transformed for each TF, with green indicating low relative activity and orange indicating high relative activity.

To explore the functional relevance of TFs at each stage, we identified 824 of the 1,442 input vertebrate TF motifs (57%) as significantly enriched in active promoter regions (adj. p < 0.05; Supplementary Table S4.5), including motifs with established roles in vertebrate development and cichlid adaptive traits. Although many motifs were enriched across all three stages, enrichment was generally strongest at 7 dpf, suggesting heightened TF activity around the embryonic-to-larval transition. Notable examples across stages included SOX10, enriched across all three stages and implicated in neural crest specification [7, 9] and divergence in cichlids and teleosts [9], and EGR1, which was most strongly enriched at 7 dpf and is associated with behavioural and social traits in cichlids [66] (Fig. 4B; Supplementary Table S4.5). Several TF motifs, including CRX, FOXA1, FOXA2, HNF4A and RXRA, were consistently enriched across all three stages but typically showed higher enrichment at 7 dpf (Fig. 4B; Supplementary Table S4.5), suggesting a conserved core regulatory programme with a mid-developmental peak. By contrast, motifs such as ELF4 and HOXC6 showed more restricted or weaker enrichment (Fig. 4B; Supplementary Table S4.5), pointing to temporal shifts in TF usage during embryogenesis.

Clustering of TF activity profiles separated motifs into groups with distinct temporal behaviour (Fig. 4C). Cluster 1 and cluster 5 showed relatively stable activity across 3, 7 and 12 dpf, consistent with a conserved developmental programme, whereas clusters 2, 3, 4 and 6 exhibited marked stage-dependent shifts (Fig. 4C), potentially indicating rewiring of TF usage during embryonic progression. The strongest stage-divergent patterns were observed in clusters 3, 4 and 6, which showed elevated activity either at 3 dpf or 7 dpf, with fewer of those cluster motifs active at 12 dpf (Fig. 4C). This suggests that TF activity is most dynamic during the transition from early *A. calliptera* embryonic patterning to later differentiation.

To distinguish broadly conserved from stage-divergent TF activity across stages, clustered motifs were subdivided into the 10 least and 10 most variable motifs, selected by row-wise standard deviation and range (Fig. 4D). The conserved group included core developmental and neurodevelopmental TFs (e.g., SOX3, TCF7 and ARX) [67], whereas the divergent group included TF activity associated with nuclear receptor and signal-responsive regulation (e.g., PPARD, RXRA and VDR) [68] (Fig. 4D). Together, these patterns suggest that stable motifs support broad embryonic programmes, while divergent modules reflect stage-specific regulatory shifts. These results are consistent with the differential accessibility analysis and support a model in which developmental progression in *A. calliptera* may be driven by both persistent core regulators and stage-specific transcriptional rewiring.

### The TF–promoter regulatory network is topologically stable across embryogenesis, with conserved developmental hub regulators

Beyond individual TF–target relationships, we examined the overall topology of the TF–promoter regulatory network inferred from footprint-supported motif enrichment at each developmental stage. Of 1,442 vertebrate and cichlid-specific TF motifs tested, a near-identical set was significantly enriched (FDR < 0.05) in accessible gene promoters at all three stages (824 at 3 dpf, 822 at 7 dpf, 824 at 12 dpf), with 822 motifs (99.8%) shared across all three and no stage-specific motifs identified (Supplementary Table 5a-b). This indicates that the compositional architecture of the promoter-level TF regulatory network, where TFs have accessible, motif-matched binding opportunities, is established early and remains largely stable across the three-stage window sampled here.

The network’s highest-connectivity (‘hub’) regulators were consistent across all three stages, led by FOXJ3, ARID3A, AR, STAT1, FOXG1, PTF1A, SOX2, SOX10, NFATC1 and STAT2 (each associated with >6,700 gene promoters per stage; Supplementary Table 5b). These hubs include well-characterised drivers of vertebrate neural and sensory development — SOX2 (neural progenitor maintenance), FOXG1 (forebrain patterning) [69], PTF1A (retinal interneuron and neural tube specification) and SOX10 (neural crest derivatives)[9] — alongside broadly expressed chromatin- and signalling-associated factors (ARID3A, STAT1/STAT2, NFATC1, AR). Rather than a developmental stage acquiring or losing major regulatory hubs, the persistence of this hub set throughout 3–12 dpf suggests a broadly conserved set of TFs operating across the 3 to 12 dpf window of *A. calliptera* development.

*A. calliptera* is considered a proxy for the generalist riverine lineage that seeded the three primary Lake Malawi lineages [70], retaining ecological and genomic characteristics, including an unspecialised habitat range and the absence of the large structural inversions fixed in several derived Lake Malawi ecogroups [71], consistent with an ancestral-like state. A topologically stable, broadly conserved TF– promoter network of this kind, dominated by hub regulators of core neural and sensory patterning, rather than stage- or lineage-restricted specialists, supports a hypothesis that retention of a flexible, largely unspecialised regulatory architecture during embryogenesis may underpin subsequent lineage-specific regulatory specialisation (e.g. of visual and craniofacial systems) as seen in derived Lake Malawi radiation members [72]. However, this will need testing using comparable data from other Lake Malawi species and of a wider window of developmental stages.

To identify candidate regulators most overrepresented among predicted target gene promoters at each stage, we ranked TFs by fold enrichment of footprint-supported motif occurrence relative to genome-wide background (Fig. 5A; Supplementary Table S5c). The strongest hub TFs differed by developmental stage: 3 dpf was dominated by zinc-finger and homeodomain factors (e.g. ZFP64, ZNF436, ZNF324, HOXB1, IRX2), whereas 7 and 12 dpf showed convergent enrichment for chromatin-organisation and cell-cycle-associated factors, including CTCF, CTCFL, ZBTB14, NRF1, TCFL5 and multiple E2F-family members, consistent with a shift from patterning-associated to proliferation- and chromatin-organisation-associated regulatory activity as development proceeds [10].

**Figure 5.**
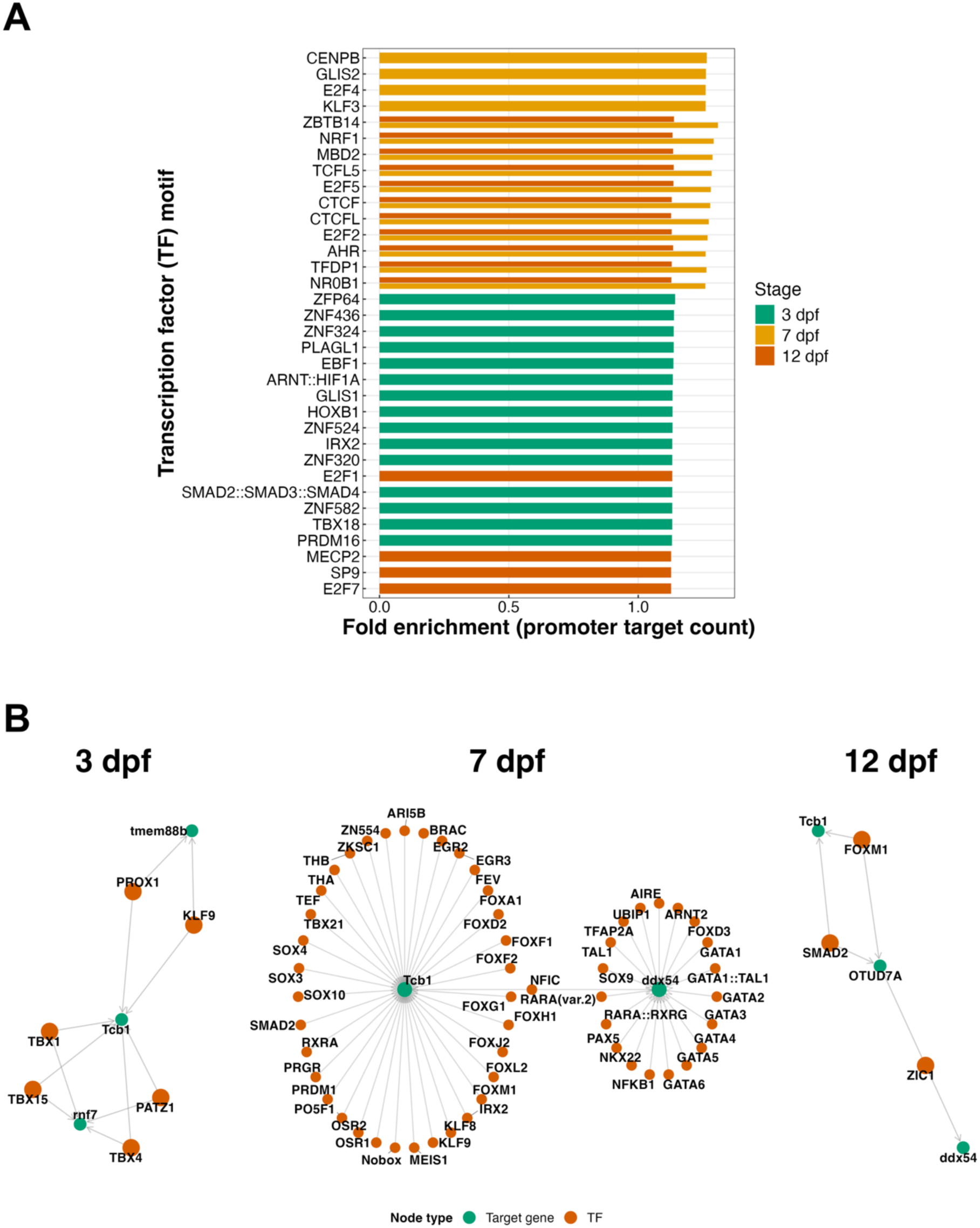
Hub TF fold enrichment and candidate TF-target gene regulatory networks across embryonic development. **(A)** Top 15 hub TFs per developmental stage, ranked by fold enrichment of footprint-supported motif occurrence among predicted target gene promoters relative to genome-wide background (Supplementary Table S5). **(B)** Candidate TF-target gene regulatory networks restricted to edges supported by accelerated conserved non-coding elements (aCNEs), shown separately for 3, 7 and 12 dpf; orange nodes are TFs and teal nodes are target genes, with edges indicating a candidate regulatory interaction.

Beyond fold enrichment, we examined the smaller set of candidate TF-target gene edges restricted to accelerated conserved non-coding elements (aCNEs) at each stage. These edges provide additional support from conserved noncoding elements identified in five cichlids (see ‘*Materials and Methods*’), but showing accelerated sequence divergence between *A. calliptera* and *O. niloticus* relative to a neutral model, and so identify candidate stage-specific regulatory interactions (Fig. 5B). At 3 dpf, the hub TFs were PROX1 and KLF9 together with the T-box factors TBX1, TBX4 and TBX15. These TFs converged on a small set of target genes, including *Tcb1*, which encodes a DNA transposase of the *Tc1/mariner* family. This family of transposable elements (TEs) is abundant in Lake Malawi cichlid genomes, including that of *A. calliptera*, and contributes substantially to their structural variation [73]. TE movement could alter regulatory sequences, including by gaining or losing transcription factor binding sites [74]. *TBX1* is also a candidate of interest: it is a conserved regulator of pharyngeal arch [75] and craniofacial patterning [76], a process that has diversified extensively in Lake Malawi cichlids. At 7 dpf, the network was subst antially denser, centred on a cluster of over 30 TFs including forkhead- (FOXA1, FOXG1, FOXH1), SOX- (SOX3, SOX10) and GATA-family (GATA1-6, GATA1::TAL1) factors together with nuclear hormone receptors (RXRA, RARA::RXRA), consistent with concurrent organogenesis, endoderm/haematopoietic differentiation and continued neural crest-derived patterning at this stage [17]. At 12 dpf, the network was again comparatively sparse, with candidate edges linking NKX6-1, NKX6-2, FOXM1, SMAD2 and ZIC1 to a small number of targets; *ZIC1* is a conserved regulator of craniofacial cartilage and somite patterning [77], and *SMAD2* mediates TGF-β/Nodal signalling in axial and organ patterning [78], making both plausible candidates for stage-specific regulatory input into late craniofacial and skeletal development. Because these aCNE-restricted networks are sparse and derived from a single species and developmental window, we present them as putative candidate interactions that could be associated with phenotypic divergence in the Lake Malawi radiation; testing individual TF-target interactions (e.g. via reporter assays or targeted perturbation [79]) could establish their functional relevance to craniofacial, neural and organ diversification across Lake Malawi cichlids more broadly.

## Discussion

Here we present the first genome-wide map of chromatin accessibility across *A. calliptera* embryonic development, spanning three key stages from the pharyngula period to early post-hatching prejuvenile development (3, 7 and 12 dpf). Accessible chromatin was pervasive throughout the genome at every stage and enriched at gene promoters and conserved noncoding elements relative to the genomic background, indicating that the core regulatory architecture of the genome is already broadly established by the earliest stage sampled here. The additional enrichment we observed at accelerated conserved noncoding elements (aCNEs) alongside highly conserved elements (hCNEs) at all three stages suggests that both deeply conserved and more rapidly evolving regulatory sequences are already active within the developmental window sampled here, consistent with the view that lineage-specific regulatory innovation may proceed through modification of existing, broadly conserved *cis-*regulatory modules [80, 81] and their accessibility.

The functional composition of genes associated with accessible promoters shifted systematically across development. Generic developmental patterning terms dominated accessible promoters at the pharyngula stage (3 dpf), the repertoire was broadest at 7 dpf, shortly after hatching (approximately stage 20), when sensory, craniofacial and musculoskeletal structures are differentiating, and neural and locomotory terms were added by 12 dpf (approximately stage 24), before yolk absorption and the onset of active feeding [17]. This progression, from broad early patterning through a diverse mid-developmental programme to a more specialised late neural/behavioural repertoire, captures the expected trajectory of vertebrate [82] and cichlid [9, 17] embryogenesis, in which early pattern-forming programmes are progressively overlaid by tissue- and system-specific differentiation programmes as organogenesis proceeds.

The sensory and retinal signals can also be related to the timing of eye development. In *A. calliptera*, the optic primordia form from the anterior neural keel at around 8–9 somites and the lenses are visible by 18 somites, during segmentation, after which retinal pigmentation begins at the pharyngula stage and intensifies until the eyes are fully opaque around hatching [17]. The 3 dpf sample (pharyngula) therefore postdates optic cup and lens formation and coincides with the onset of retinal pigmentation, consistent with retinal and sensory terms already being significant at this stage, whereas 7 dpf follows hatching, when the eyes are fully pigmented, and terms linked to retinal differentiation (neural retina development, photoreceptor cell differentiation) were specific to this stage. That promoters at neural and retinal genes that opened between 3 and 7 dpf (*neurod4, irx7, notch1b, isl2b*) were closed again by 12 dpf suggests a window of retinal specification and differentiation spanning the pharyngula to early post-hatching period, although our whole-embryo design cannot resolve tissue-specific timing, and a direct comparison with the timing of retinal neurogenesis and photoreceptor differentiation will require tissue-resolved data.

Differential accessibility analysis provided a dynamic view of this process: across all three pairwise comparisons, and particularly the two comparisons involving 3 dpf, promoter accessibility change was biased towards gains rather than losses of accessibility. Genes gaining accessibility during the earliest transitions were enriched almost exclusively for broad metabolic and biosynthetic processes rather than specific developmental terms, consistent with a general, genome-wide increase in transcriptional and translational processes as embryos progress through early growth and tissue differentiation. The smaller set of changes between 7 and 12 dpf was more specific: gains were enriched for apoptotic and cytoskeletal remodelling terms, consistent with the role of programmed cell death and tissue remodelling in later embryogenesis [83], while losses were enriched for neural plate and retinal patterning terms, which may reflect early neural patterning becoming less accessible once it is complete. Notably, almost all late losses occurred at promoters that had opened between 3 and 7 dpf, indicating transient rather than sustained opening during a window of neural and retinal specification. This progression from broad to more targeted regulatory change is in line with other vertebrates, where evolutionarily newer regulatory regions become accessible later in embryogenesis, after the phylotypic period [84], and with the stage-specific expression of neural, retinal and craniofacial programmes in Lake Malawi cichlids [7]. To our knowledge, chromatin accessibility has not previously been profiled across cichlid embryogenesis, and tissue attributions here are inferred from GO enrichment of whole-embryo data only. Together, these results suggest that the dominant chromatin-level signature of early *A. calliptera* embryogenesis is one of broad regulatory opening in support of general cellular activity, with more targeted, direction-specific remodelling emerging only later as specific tissues mature.

Transcription factor footprinting identified stage-specific activity among developmentally relevant regulators, while analysis of the broader TF–promoter network topology showed that the underlying set of TFs with accessible, motif-matched promoter binding opportunities is stable across the same developmental window. This is dominated by hub regulators of core neural and sensory patterning (e.g. SOX2, FOXG1, PTF1A, SOX10). Taken together with the differential accessibility patterns identified, this suggests a model in which the compositional “wiring” of the promoter-level regulatory network is established early and largely maintained, while developmental progression is instead achieved through changes in the activity and accessibility of a broadly conserved set of regulatory elements and regulators, rather than early rewiring of which factors can act at each stage. The stability of this network from the pharyngula stage onwards is compatible with the hourglass model of embryogenesis, in which the mid-embryonic phylotypic period is the most conserved and earlier and later stages diverge more [82, 85]; testing this would, however, require comparison with earlier (cleavage and gastrula) stages and with other species, which we have not sampled. Testing along a wider developmental window would be useful to also assess any interspecies and inter-tissue rewiring of TFs, and whether the developmental core established here may itself become a substrate for lineage- and tissue-specific rewiring later in cichlid life history [6, 38, 44].

*A. calliptera* is considered a proxy for the generalist riverine lineage that seeded the explosive radiation of Lake Malawi lineages [70], retaining an unspecialised ecology and genomic characteristics [71]. The broadly conserved, flexible regulatory architecture characterised here — a stable core TF–promoter network, widespread rather than narrowly targeted chromatin opening during early development, and enrichment of both deeply conserved and more rapidly evolving noncoding elements — is compatible with the hypothesis that retention of such a flexible embryonic regulatory baseline may have been permissive for the subsequent lineage-specific regulatory specialisation documented in derived Lake Malawi cichlids, for example in visual and craniofacial systems [6, 33, 86]. This developmental epigenomic resource therefore complements existing genomic [5, 8, 73, 87, 88] and adult-tissue epigenomic [6, 89] studies of cichlid diversification, by providing an embryonic baseline against which regulatory changes in more derived, ecologically specialised lineages can ultimately be compared.

This study however has a few limitations. Firstly, with two replicates per stage, statistical power to detect more subtle accessibility changes, particularly for genes with small effect sizes, remains limited. Secondly, ATAC-seq was performed on whole embryos rather than single cells, so the signals reported here represent an average across all cell types present at each stage and likely masks cell-type-specific regulatory dynamics, particularly for less abundant cell populations [90]. Finally, we were unable to sequence the transcriptome from corresponding embryos and thus, could not test whether accessibility correlated with expression changes of the predicted target genes. Future work should therefore extend this dataset to additional developmental stages, incorporating joint single-cell or nuclei chromatin and transcriptomic profiling [90], with directly comparing embryonic regulatory landscapes between *A. calliptera* and more ecologically specialised Lake Malawi cichlids [91]. This would help test whether the flexible, broadly conserved regulatory architecture documented here is indeed a shared ancestral feature that has been differentially specialised during the radiation, or a lineage-specific feature of *A. calliptera* itself.

## Conclusions

We generated the first genome-wide, developmental-stage-resolved map of chromatin accessibility in *Astatotilapia calliptera*, a generalist lineage considered representative of the ancestral riverine stock that seeded the Lake Malawi cichlid radiation. Accessible chromatin was pervasive and enriched at conserved regulatory elements throughout embryogenesis, with the functional character of accessible promoters shifting from broad developmental patterning at 3 dpf to a diverse, then increasingly neural and behavioural, gene repertoire by 12 dpf. Differential accessibility analysis showed that chromatin opening, rather than closing, dominates the earliest developmental transitions, and transcription factor footprinting revealed a topologically stable promoter-regulatory network built around conserved developmental hub regulators. Together, these findings establish a developmental epigenomic baseline for a key generalist lineage in the cichlid adaptive radiation and provide a framework for future comparative studies of how regulatory evolution has contributed to the remarkable phenotypic diversification of Lake Malawi cichlid fishes.

## Declarations

### Ethics approval and consent to participate

All animal procedures were approved by the University of Hull Animal Welfare and Ethical Review Body (AWERB) and carried out in accordance with approved guidelines.

### Consent for publication

Not applicable.

### Availability of data and materials

The data underlying this article are available in the article, its online supplementary material, or uploaded to Zenodo (https://doi.org/10.5281/zenodo.23258231). All code is available on GitHub at: https://github.com/tmehta12/Acalliptera_ATAC.

### Authors’ contributions

TKM: conceptualisation, methodology, transcription factor footprinting and regulatory network analysis, supervision, writing – original draft, review and editing. M: peak annotation, functional enrichment and differential accessibility analysis, writing – review and editing. AMS, DJ: resources (embryo collection, tissue dissection), writing – review and editing. FD-P, WH: funding acquisition, supervision, writing – review and editing. All authors read and approved the final manuscript.

## Acknowledgements

The authors acknowledge the support of the Biotechnology and Biological Sciences Research Council (BBSRC), part of UK Research and Innovation. The authors would like to acknowledge Tom Barker, Vanda Knitlhoffer, Leah Catchpole, and Suzanne Henderson of the Genomics Pipelines Group at Earlham Institute for data generation including preparation of library pooling, and sequencing. The authors would also like to acknowledge the Scientific Computing group, as well as support for the physical HPC infrastructure and data centre delivered via the NBI Research Computing group.

## References

1. Kocher TD: Adaptive evolution and explosive speciation: the cichlid fish model. Nature Reviews Genetics 2004, 5(4):288–298.

2. Malinsky M, Svardal H, Tyers AM, Miska EA, Genner MJ, Turner GF, Durbin R: Whole-genome sequences of Malawi cichlids reveal multiple radiations interconnected by gene flow. Nature ecology & evolution 2018, 2(12):1940–1955.

3. Salzburger W: Understanding explosive diversification through cichlid fish genomics. Nature Reviews Genetics 2018, 19(11):705–717.

4. Brawand D, Wagner CE, Li YI, Malinsky M, Keller I, Fan S, Simakov O, Ng AY, Lim ZW, Bezault E: The genomic substrate for adaptive radiation in African cichlid fish. Nature 2014, 513(7518):375–381.

5. Svardal H, Salzburger W, Malinsky M: Genetic Variation and Hybridization in Evolutionary Radiations of Cichlid Fishes. Annu Rev Anim Biosci 2021, 9:55–79.

6. Mehta TK, Man AL, Etherington G, Smith AM, Indermaur A, Salzburger W, Joyce D, Di-Palma F, Haerty W: Genome-wide evolution of lineage-specific chromatin accessibility links regulatory changes to cichlid phenotypic diversity. Genome Biol Evol 2026.

7. Bloomquist RF, Fowler TE, Sylvester JB, Miro RJ, Streelman JT: A compendium of developmental gene expression in Lake Malawi cichlid fishes. BMC Developmental Biology 2017, 17(1):3–3.

8. Svardal H, Quah FX, Malinsky M, Ngatunga BP, Miska EA, Salzburger W, Genner MJ, Turner GF, Durbin R: Ancestral hybridization facilitated species diversification in the Lake Malawi cichlid fish adaptive radiation. Molecular biology and evolution 2020, 37(4):1100–1113.

9. Marconi A, Vernaz G, Karunaratna A, Ngochera MJ, Durbin R, Santos ME: Genetic and Developmental Divergence in the Neural Crest Program between Cichlid Fish Species. Mol Biol Evol 2024, 41(11).

10. Louise Smith E, Mok GF, Münsterberg A: Investigating chromatin accessibility during development and differentiation by ATAC-sequencing to guide the identification of cis-regulatory elements. Biochemical Society Transactions 2022, 50(3):1167–1177.

11. Bentsen M, Goymann P, Schultheis H, Klee K, Petrova A, Wiegandt R, Fust A, Preussner J, Kuenne C, Braun T: ATAC-seq footprinting unravels kinetics of transcription factor binding during zygotic genome activation. Nature communications 2020, 11(1):4267.

12. Buenrostro JD, Giresi PG, Zaba LC, Chang HY, Greenleaf WJ: Transposition of native chromatin for fast and sensitive epigenomic profiling of open chromatin, DNA-binding proteins and nucleosome position. Nature methods 2013, 10(12):1213–1218.

13. Li Z, Schulz MH, Look T, Begemann M, Zenke M, Costa IG: Identification of transcription factor binding sites using ATAC-seq. Genome biology 2019, 20(1):45.

14. Mehta TK, Man A, Ciezarek A, Ranson K, Penman D, Di-Palma F, Haerty W: Chromatin accessibility in gill tissue identifies candidate genes and loci associated with aquaculture relevant traits in tilapia. Genomics 2023, 115(4):110633.

15. Yuan Y, Xi H, Chen S, Zhou Z, Zhu W, Liu Y, Li Y, Shen Z, Wang C, Xiao J: Integration of ATAC-seq and RNA-seq unravels chromatin accessibility associated with gonad development after sex reversal in GIFT tilapia. Genomics 2026:111230.

16. Bogdanović O, Fernandez-Miñán A, Tena JJ, Calle-Mustienes Edl, Hidalgo C, Kruysbergen Iv, Heeringen SJv, Veenstra GJC, Gómez-Skarmeta JL: Dynamics of enhancer chromatin signatures mark the transition from pluripotency to cell specification during embryogenesis. Genome Research 2012, 22(10).

17. Marconi A, Yang CZ, McKay S, Santos ME: Morphological and temporal variation in early embryogenesis contributes to species divergence in Malawi cichlid fishes. Evol Dev 2023, 25(2):170–193.

18. Andrews S: FastQC: A quality control tool for high throughput sequence data [Online]. *Available online at:* http://wwwbioinformaticsbabrahamacuk/projects/fastqc/ 2010.

19. Langmead B, Salzberg SL: Fast gapped-read alignment with Bowtie 2. In: Nature Methods. vol. 9; 2012: 357–359.

20. Danecek P, Bonfield JK, Liddle J, Marshall J, Ohan V, Pollard MO, Whitwham A, Keane T, McCarthy SA, Davies RM et al: Twelve years of SAMtools and BCFtools. GigaScience 2021, 10(2):giab008–giab008.

21. Camacho C, Coulouris G, Avagyan V, Ma N, Papadopoulos J, Bealer K, Madden TL: BLAST+: Architecture and applications. BMC Bioinformatics 2009, 10:421–421.

22. Tarasov A, Vilella AJ, Cuppen E, Nijman IJ, Prins P: Sambamba: fast processing of NGS alignment formats. Bioinformatics 2015, 31(12):2032– 2034.

23. Quinlan AR, Hall IM: BEDTools: A flexible suite of utilities for comparing genomic features. Bioinformatics 2010, 26(6):841–842.

24. Feng J, Liu T, Qin B, Zhang Y, Liu XS: Identifying ChIP-seq enrichment using MACS. Nature Protocols 2012, 7(9):1728–1740.

25. Zhang Y, Liu T, Meyer CA, Eeckhoute J, Johnson DS, Bernstein BE, Nusbaum C, Myers RM, Brown M, Li W et al: Model-based analysis of ChIP-Seq (MACS). Genome biology 2008, 9(9):R137–R137.

26. Ou J, Liu H, Yu J, Kelliher MA, Castilla LH, Lawson ND, Zhu LJ: ATACseqQC: a Bioconductor package for post-alignment quality assessment of ATAC-seq data. BMC Genomics 2018, 19(1):169–169.

27. Benjamini Y, Hochberg Y: Benjamini Y, Hochberg Y. Controlling the false discovery rate: a practical and powerful approach to multiple testing. Journal of the Royal Statistical Society B 1995, 57(1):289–300.

28. Kolberg L, Raudvere U, Kuzmin I, Adler P, Vilo J, Peterson H: g:Profiler-interoperable web service for functional enrichment analysis and gene identifier mapping (2023 update). Nucleic Acids Res 2023, 51(W1):W207– W212.

29. Lun AT, Smyth GK: csaw: a Bioconductor package for differential binding analysis of ChIP-seq data using sliding windows. Nucleic Acids Res 2016, 44(5):e45.

30. Chen Y, Chen L, Lun ATL, Baldoni PL, Smyth GK: edgeR v4: powerful differential analysis of sequencing data with expanded functionality and improved support for small counts and larger datasets. Nucleic Acids Res 2025, 53(2).

31. Conte Ma, Kocher TD: An improved genome reference for the African cichlid, Metriaclima zebra. BMC Genomics 2015, 16(1):724–724.

32. Conte MA, Joshi R, Moore EC, Nandamuri SP, Gammerdinger WJ, Roberts RB, Carleton KL, Lien S, Kocher TD: Chromosome-scale assemblies reveal the structural evolution of African cichlid genomes. GigaScience 2019, 8(4):giz030–giz030.

33. Brawand D, Wagner CE, Li YI, Malinsky M, Keller I, Fan S, Simakov O, Ng aY, Lim ZW, Bezault E et al: The genomic substrate for adaptive radiation in African cichlid fish. Nature 2014, 2(c):17.

34. Conte MA, Gammerdinger WJ, Bartie KL, Penman DJ, Kocher TD: A high quality assembly of the Nile Tilapia (Oreochromis niloticus) genome reveals the structure of two sex determination regions. BMC Genomics 2017, 18(1):341–341.

35. Armstrong J, Hickey G, Diekhans M, Fiddes IT, Novak AM, Deran A, Fang Q, Xie D, Feng S, Stiller J et al: Progressive Cactus is a multiple-genome aligner for the thousand-genome era. Nature 2020, 587(7833):246–251.

36. Hubisz MJ, Pollard KS, Siepel A: PHAST and RPHAST: phylogenetic analysis with space/time models. Briefings in Bioinformatics 2011, 12(1):41–51.

37. Pollard K, Hubisz MJ, Rosenbloom KR, Siepel A: Detection of nonneutral substitution rates on mammalian phylogenies. Genome Res 2010, 20(1):110–121.

38. Mehta T, Koch C, Nash W, Knaack SA, Sudhakar P, Olbei M, Bastkowski S, Penso-Dolfin L, Korcsmaros T, Haerty W et al: Evolution of regulatory networks associated with traits under selection in cichlids. Genome Biol 2021, 22(1):25–25.

39. Castro-Mondragon JA, Riudavets-Puig R, Rauluseviciute I, Berhanu Lemma R, Turchi L, Blanc-Mathieu R, Lucas J, Boddie P, Khan A, Manosalva Pérez N et al: JASPAR 2022: the 9th release of the open-access database of transcription factor binding profiles. Nucleic Acids Research 2022, 50(D1):D165–D173.

40. Rauluseviciute I, Riudavets-Puig R, Blanc-Mathieu R, Castro-Mondragon JA, Ferenc K, Kumar V, Lemma RB, Lucas J, Cheneby J, Baranasic D et al: JASPAR 2024: 20th anniversary of the open-access database of transcription factor binding profiles. Nucleic Acids Res 2024, 52(D1):D174–D182.

41. Kulakovskiy IV, Medvedeva YA, Schaefer U, Kasianov AS, Vorontsov IE, Bajic VB, Makeev VJ: HOCOMOCO: A comprehensive collection of human transcription factor binding sites models. Nucleic Acids Research 2013, 41(D1):D195–202.

42. Heger A, Webber C, Goodson M, Ponting CP, Lunter G: GAT: A simulation framework for testing the association of genomic intervals. Bioinformatics 2013, 29(16):2046–2048.

43. Li Z, Schulz MH, Look T, Begemann M, Zenke M, Costa IG: Identification of transcription factor binding sites using ATAC-seq. Genome Biology 2019, 20(1):45–45.

44. Mehta T, Penso-Dolfin L, Nash W, Roy S, Di-Palma F, Haerty W: Evolution of miRNA-binding sites and regulatory networks in cichlids. MBE 2022, 39(7):146.

45. Buchmann K, Karami AM, Duan Y: The early ontogenetic development of immune cells and organs in teleosts. Fish Shellfish Immunol 2024, 146:109371.

46. Tittle RK, Sze R, Ng A, Nuckels RJ, Swartz ME, Anderson RM, Bosch J, Stainier DY, Eberhart JK, Gross JM: Uhrf1 and Dnmt1 are required for development and maintenance of the zebrafish lens. Dev Biol 2011, 350(1):50–63.

47. Angileri KM, Gross JM: dnmt1 function is required to maintain retinal stem cells within the ciliary marginal zone of the zebrafish eye. Sci Rep 2020, 10(1):11293.

48. Lane BM, Lister JA: Otx but Not Mitf Transcription Factors Are Required for Zebrafish Retinal Pigment Epithelium Development. PLoS ONE 2012, 7(11).

49. Sun K, Liu X, Xu R, Liu C, Meng A, Lan X: Mapping the chromatin accessibility landscape of zebrafish embryogenesis at single-cell resolution by SPATAC-seq. Nat Cell Biol 2024, 26(7):1187–1199.

50. Becker EA, Bird NC, Webb JF: Post-embryonic development of canal and superficial neuromasts and the generation of two cranial lateral line phenotypes. J Morphol 2016, 277(10):1273–1291.

51. Kamler E: Resource allocation in yolk-feeding fish. Reviews in Fish Biology and Fisheries 2008, 18(2):143–200.

52. Schmidt R, Strähle U, Scholpp S: Neurogenesis in zebrafish - from embryo to adult. Neural Dev 2013, 8:3.

53. Inoue T, Hojo M, Bessho Y, Tano Y, Lee JE, Kageyama R: Math3 and NeuroD regulate amacrine cell fate specification in the retina. Development 2002, 129(4):831–842.

54. Zhang Y, Yang Y, Trujillo C, Zhong W, Leung YF: The expression of irx7 in the inner nuclear layer of zebrafish retina is essential for a proper retinal development and lamination. PLoS One 2012, 7(4):e36145.

55. Pittman AJ, Law MY, Chien CB: Pathfinding in a large vertebrate axon tract: isotypic interactions guide retinotectal axons at multiple choice points. Development 2008, 135(17):2865–2871.

56. Sandkam BA, Campello L, O’Brien C, Nandamuri SP, Gammerdinger W, Conte M, Swaroop A, Carleton KL: Tbx2a modulates switching of RH2 and LWS opsin gene expression. Molecular Biology and Evolution 2020, 37(7):2002–2014.

57. Niwa R, Nagata-Ohashi K, Takeichi M, Mizuno K, Uemura T: Control of actin reorganization by Slingshot, a family of phosphatases that dephosphorylate ADF/cofilin. Cell 2002, 108(2):233–246.

58. Jia S, Nakaya N, Piatigorsky J: Differential expression patterns and developmental roles of duplicated scinderin-like genes in zebrafish. Developmental Dynamics 2009, 238(10):2633–2640.

59. Sidhaye J, Pinto CS, Dharap S, Jacob T, Bhargava S, Sonawane M: The zebrafish goosepimples/myosin Vb mutant exhibits cellular attributes of human microvillus inclusion disease. Mech Dev 2016, 142:62–74.

60. Jukam D, Shariati SAM, Skotheim JM: Zygotic Genome Activation in Vertebrates. Dev Cell 2017, 42(4):316–332.

61. Coyle MC, King N: The evolutionary foundations of transcriptional regulation in animals. Nat Rev Genet 2025, 26(12):812–827.

62. Vierstra J, Lazar J, Sandstrom R, Halow J, Lee K, Bates D, Diegel M, Dunn D, Neri F, Haugen E et al: Global reference mapping of human transcription factor footprints. Nature 2020, 583(7818):729–736.

63. Macdonald R, Barth KA, Xu Q, Holder N, Mikkola I, Wilson SW: Midline signalling is required for Pax gene regulation and patterning of the eyes. Development 1995, 121(10):3267–3278.

64. Lee CS, Friedman JR, Fulmer JT, Kaestner KH: The initiation of liver development is dependent on Foxa transcription factors. Nature 2005, 435(7044):944–947.

65. Dee CT, Szymoniuk CR, Mills PE, Takahashi T: Defective neural crest migration revealed by a Zebrafish model of Alx1-related frontonasal dysplasia. Hum Mol Genet 2013, 22(2):239–251.

66. Burmeister SS, Jarvis ED, Fernald RD: Rapid behavioral and genomic responses to social opportunity. PLoS Biology 2005, 3(11):1996–2004.

67. Stevanovic M, Drakulic D, Lazic A, Ninkovic DS, Schwirtlich M, Mojsin M: SOX Transcription Factors as Important Regulators of Neuronal and Glial Differentiation During Nervous System Development and Adult Neurogenesis. Front Mol Neurosci 2021, 14:654031.

68. Chung AC, Cooney AJ: The varied roles of nuclear receptors during vertebrate embryonic development. Nucl Recept Signal 2003, 1:e007.

69. Sylvester JB, Rich CA, Yi C, Peres JN, Houart C, Streelman JT: Competing signals drive telencephalon diversity. Nature Communications 2013, 4:1745–1745.

70. Parsons PJ, Bridle JR, Ruber L, Genner MJ: Evolutionary divergence in life history traits among populations of the Lake Malawi cichlid fish Astatotilapia calliptera. Ecol Evol 2017, 7(20):8488–8506.

71. Kumar NM, Cooper TL, Kocher TD, Streelman JT, McGrath PT: Large inversions in Lake Malawi cichlids are associated with habitat preference, lineage, and sex determination. In.: eLife Sciences Publications, Ltd; 2025.

72. Salzburger W: Understanding explosive diversification through cichlid fish genomics. Nature Rev Genet 2018, 19:705–717–705–717.

73. Quah FX, Almeida MV, Blumer M, Yuan CU, Fischer B, See K, Jackson B, Zatha R, Rusuwa B, Turner GF et al: Lake Malawi cichlid pangenome graph reveals extensive structural variation driven by transposable elements. Genome Res 2025, 35(5):1094–1107.

74. Carleton KL, Conte MA, Malinsky M, Nandamuri SP, Sandkam BA, Meier JI, Mwaiko S, Seehausen O, Kocher TD: Movement of transposable elements contributes to cichlid diversity. Mol Ecol 2020, 29(24):4956–4969.

75. Choe CP, Crump JG: Tbx1 controls the morphogenesis of pharyngeal pouch epithelia through mesodermal Wnt11r and Fgf8a. Development 2014, 141(18):3583–3593.

76. Albertson RC, Kocher TD: Genetic and developmental basis of cichlid trophic diversity. Heredity 2006, 97(3):211–221.

77. Houtmeyers R, Souopgui J, Tejpar S, Arkell R: The ZIC gene family encodes multi-functional proteins essential for patterning and morphogenesis. Cell Mol Life Sci 2013, 70(20):3791–3811.

78. Jia S, Ren Z, Li X, Zheng Y, Meng A: smad2 and smad3 are required for mesendoderm induction by transforming growth factor-beta/nodal signals in zebrafish. J Biol Chem 2008, 283(4):2418–2426.

79. Clark B, Kuwalekar M, Fischer B, Woltering J, Biran J, Juntti S, Kratochwil CF, Santos ME, Almeida MV: Genome editing in East African cichlids and tilapias: state-of-the-art and future directions. Open Biol 2023, 13(11):230257.

80. Wray GA: The evolutionary significance of cis-regulatory mutations. In: Nature Reviews Genetics. vol. 8; 2007: 206–216.

81. Pauls S, Goode DK, Petrone L, Oliveri P, Elgar G: Evolution of lineage-specific functions in ancient cis-regulatory modules. Open Biol 2015, 5(11).

82. Irie N, Kuratani S: Comparative transcriptome analysis reveals vertebrate phylotypic period during organogenesis. Nat Commun 2011, 2:248.

83. Fuchs Y, Steller H: Programmed cell death in animal development and disease. Cell 2011, 147(4):742–758.

84. Uesaka M, Kuratani S, Takeda H, Irie N: Recapitulation-like developmental transitions of chromatin accessibility in vertebrates. Zoological Letters 2019, 5(1):33–33.

85. Irie N, Kuratani S: The developmental hourglass model: a predictor of the basic body plan? Development 2014, 141(24):4649–4655.

86. Patil C, Sylvester JB, Abdilleh K, Norsworthy MW, Pottin K, Malinsky M, Bloomquist RF, Johnson ZV, McGrath PT, Streelman JT: Genome-enabled discovery of evolutionary divergence in brains and behavior. Sci Rep 2021, 11(1):13016.

87. Malinsky M, Svardal H, Tyers AM, Miska EA, Genner MJ, Turner GF, Durbin R: Whole-genome sequences of Malawi cichlids reveal multiple radiations interconnected by gene flow. Nature Ecol & Evol 2018, 2(12):1940–1955.

88. Blumer LM, Burskaia V, Artiushin I, Saha J, Garcia JC, Jimenez FC, Hooft van der Huysdynen A, Elkin J, Fischer B, Van Houtte N et al: Introgression dynamics of sex-linked chromosomal inversions shape the Malawi cichlid radiation. Science 2025, 388(6752):eadr9961.

89. Vernaz G, Malinsky M, Svardal H, Du M, Tyers AM, Santos ME, Durbin R, Genner MJ, Turner GF, Miska EA: Mapping epigenetic divergence in the massive radiation of Lake Malawi cichlid fishes. Nature Communications 2021, 12(1):5870–5870.

90. Liu C, Li X, Hu Q, Jia Z, Ye Q, Wang X, Zhao K, Liu L, Wang M: Decoding the blueprints of embryo development with single-cell and spatial omics. Semin Cell Dev Biol 2025, 167:22–39.

91. Santos ME, Lopes JF, Kratochwil CF: East African cichlid fishes. Evodevo 2023, 14(1):1.

