## Supplementary Figures for "Genome-wide chromatin accessibility during early embryogenesis in a generalist cichlid fish"

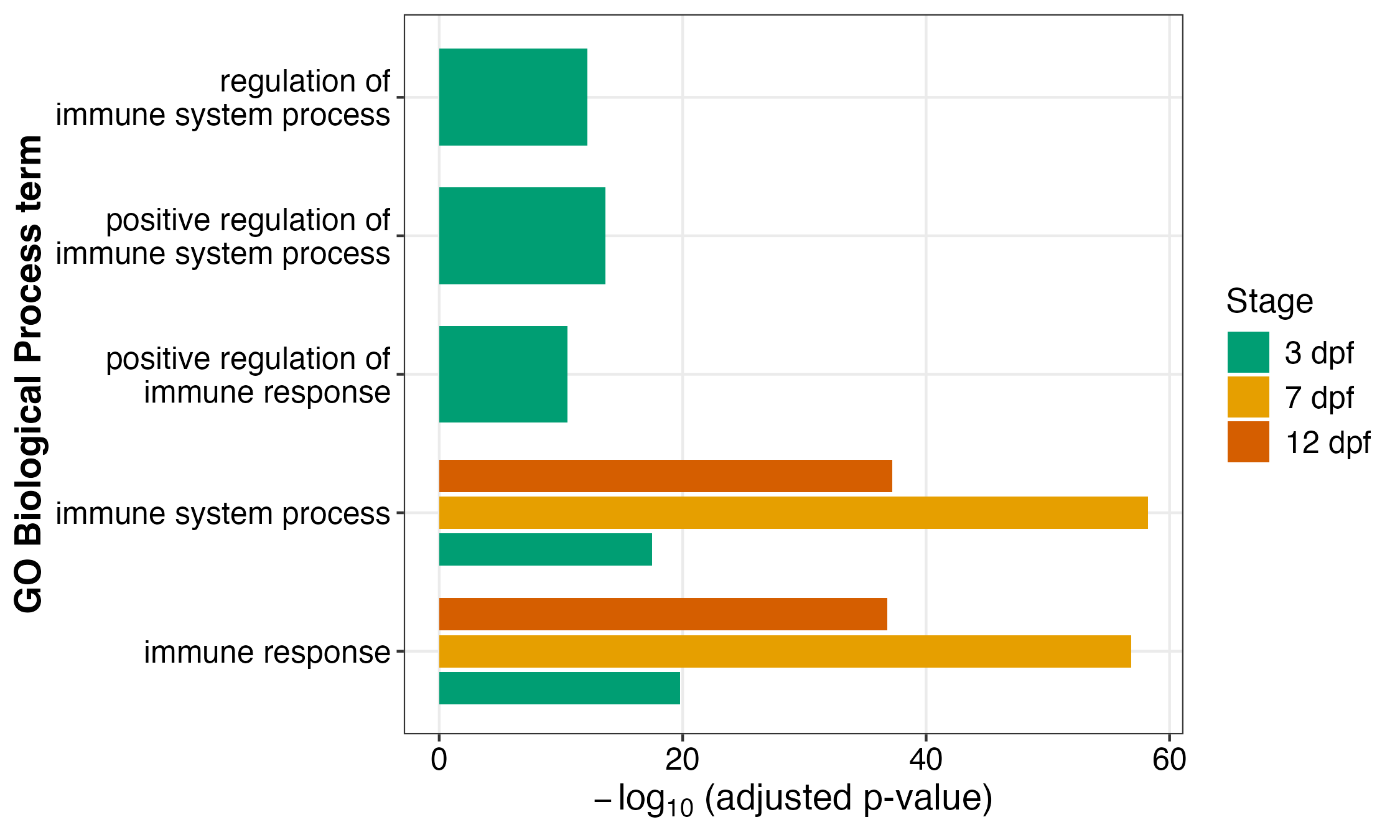

**Supplementary Figure 1.** Gene Ontology (GO) Biological Process enrichment of genes with no promoter-overlapping accessible peak at each developmental stage (specificity control). Bars show -log10(adjusted p-value) for immune system process-related GO:BP terms among genes lacking a promoter-overlapping accessible peak at 3, 7 and 12 dpf (6,613, 8,097 and 5,869 genes respectively, tested against the same background as the main promoter-accessible gene set); all terms shown were significant at adjusted p < 10⁻²⁰ at every stage. Unlike genes with accessible promoters (Figure 2), these genes were not enriched for any developmental term, supporting the specificity of the developmental GO enrichment reported for promoter-accessible genes.

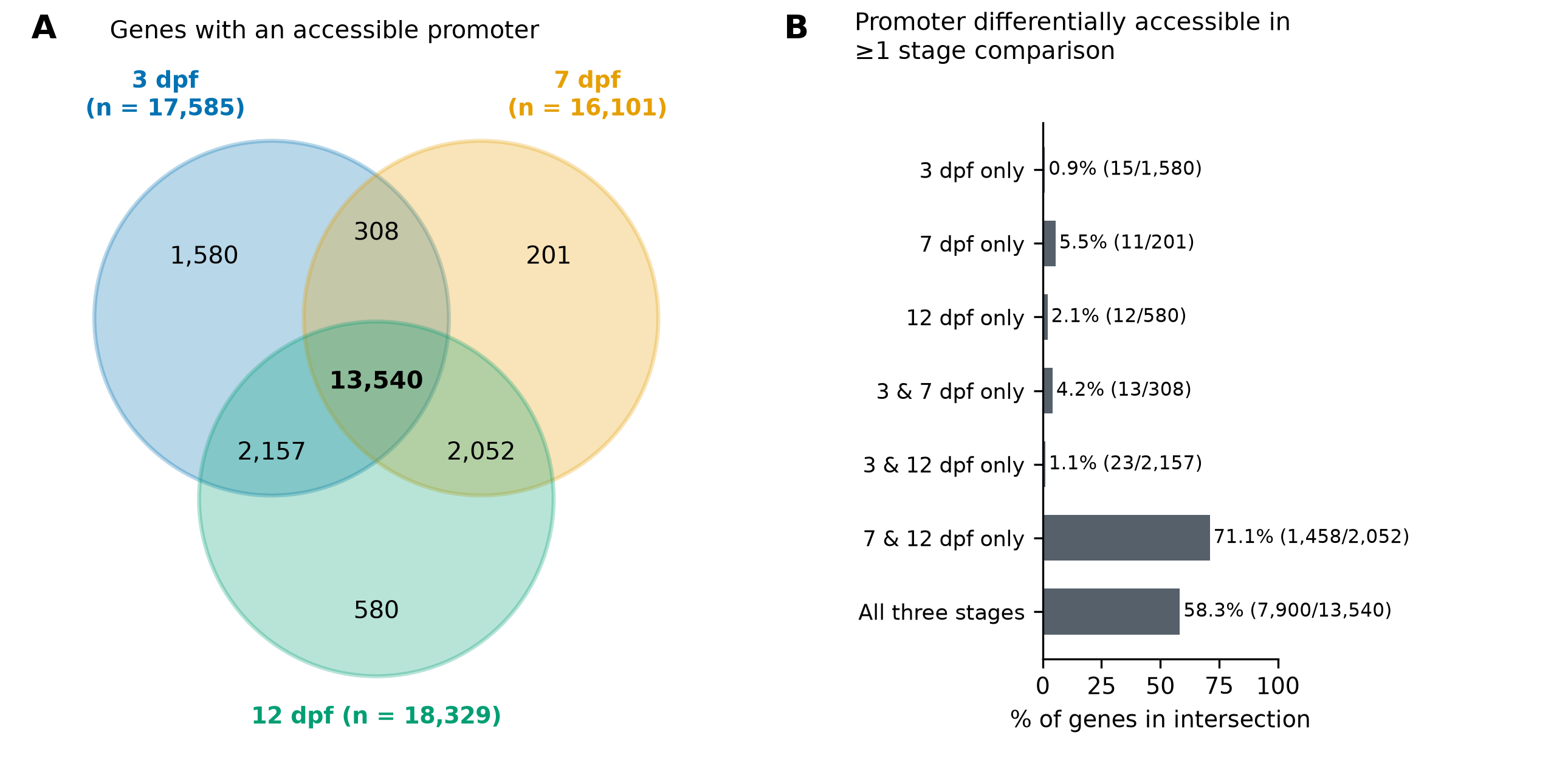

**Supplementary Figure 2.** Overlap of genes with accessible promoters across developmental stages, and their differential promoter accessibility. **(A)** Venn diagram of genes with at least one accessible peak overlapping the 5 kb promoter region at 3 dpf (n = 17,585), 7 dpf (n = 16,101) and 12 dpf (n = 18,329). Peaks are the pooled calls from the two biological replicates of each stage. In total, 20,418 of the 24,198 annotated genes have an accessible promoter at one or more stages, and 13,540 have one at all three stages. **(B)** Percentage of genes in each intersection in (A) whose promoter is differentially accessible (csaw/edgeR; region-level FDR < 0.05) in at least one of the three pairwise stage comparisons (3 vs 7, 3 vs 12 and 7 vs 12 dpf). Numbers in brackets give differentially accessible genes / genes in the intersection. Gene lists per intersection are provided in Supplementary Table S1G.
